# Species delimitation of eight ascidian (Tunicata) species from the North Eastern Mediterranean

**DOI:** 10.1101/2023.01.17.523747

**Authors:** Arzu Karahan, Berivan Temiz, Esra Öztürk, Jacob Douek, Baruch Rinkevich

**Affiliations:** Middle East Technical University, Institute of Marine Science, Department of Marine Biology and Fisheries, Mersin, Turkiye; Developmental Biology and Genomics Laboratory, Department of Anatomy, Otago School of Medical Sciences, University of Otago, P.O. Box 56, Dunedin 9054, New Zealand; Israel Oceanographic and Limnology Research, National Institute of Oceanography, Tel Shikmona, PO Box 9753, Haifa 3109701, Israel

**Author notes:** Corresponding author: Arzu Karahan, Middle East Technical University, Institute of Marine Sciences, Erdemli, Mersin, Turkiye.

**Keywords:** Botryllid ascidians, *Didemnum Polyclinum*, *Symplegma*, COI, species delimitation, Turkiye, Mediterranean Sea

## Abstract

Members of the tunicates, a subphylum of marine filter-feeder chordates, inhabit all marine and oceanic habitats from the subtidal to the abyssal. Considered as the closest relatives to the vertebrates, the tunicates are widely used as model organisms for evo-devo, allorecognition, senescence, and whole-body regeneration studies. Yet, species boundaries are poorly understood due to high morphological and genetic plasticity that characterize many tunicates taxa. Here we study taxonomy and the distribution of eight tunicate species (*Botrylloides niger Herdman, 1886/ aff. leachii, Botrylloides israeliense* Brunetti, 2009, *Botrylloides sp., Botrylloides anceps* (Herdman, 1891), *Botryllus schlosseri* (Pallas, 1766), *Didemnum perlucidum* Monniot F., 1983, *Symplegma brakenhielmi* (Michaelsen, 1904) and *Polyclinum constellatum* Savigny, 1816) sampled from six Turkish North Eastern Mediterranean Sea sites and employed the mitochondrial barcoding marker (COI) for evaluating the relationships among geographically restricted and widely spread ascidian species. Species delimitation analysis was conducted using NCBI and the present study sequences. Morphological examinations were first done in the field and then, styelide colonies were cultured in the laboratory and studied using stereo and light microscopes. A putative new *Botrylloides* species (*Botrylloides sp*.) from the Antalya region was revealed, with 99% matching on the COI gene from Saudi Arabia, further awaiting for detailed traditional taxonomy.

## Introduction

The ascidians (Phylum: Chordata, Subphylum; Tunicata) are a class of marine filter feeder organisms with ca. 3000 described species (https://www.marinespecies.org/ascidiacea) that inhabit all marine and oceanic habitats from the subtidal to the abyssal zone. As the closest relatives of the vertebrates, the tunicates are widely used as model organisms for evo-devo research analyses, for elucidating the evolution of immunity, senescence and ageing processes, for whole-body regeneration phenomena, stem cell biology, and more. Yet, species boundaries for many clades are poorly understood due to high genetic and morphological plasticity, revealing high cases of cryptic diversity (Burnet, 1971; Rinkevich *et al*., 1993; Denoeud *et al*., 2010; Reem *et al*., 2017, 2022; Viard *et al*., 2019). Further, ascidians’ classical classification requires experienced taxonomists (Rubinstein *et al*., 2013), a vanishing scientific discipline, in addition to the difficulties in assigning differentiating taxonomic characteristics between closely related species (Rocha *et al*., 2012). Nowadays, in addition to classical taxonomy, researchers use a wide range of molecular tools to elucidate biodiversity and to solve emerging species delineation issues (Karahan *et al*., 2016; Brunetti *et al*., 2017; Viard *et al*., 2019; Reem *et al*., 2017, 2022). The cytochrome oxidase subunit 1 (COI) gene is one of the most commonly used markers for this purpose (Hebert *et al*., 2003).

Studies on the taxonomy and distribution of Mediterranean Sea ascidians, now almost bicentennial old (Schlosser and Ellis, 1756; Spallanzani & Chiereghin, 1784; Savigny, 1816) were concentrated primarily on western European coasts and seas, neglecting the eastern basin areas (Berrill, 1950; Rinkevich *et al*., 1993). One example is the Levant, an area experiencing a continuous flow of non-indigenous species (NIS), including new exotic tropical ascidian species, entering the Mediterranean Sea through the Suez Canal (López-Legentil *et al*., 2015; Zenetos *et al*., 2017; Galil *et al*., 2018). Another less studied area is the Turkish Mediterranean coastline, where the knowledge on ascidians has improved considerably in the past vicenary. So far, about 50 ascidian species (native and NIS), consisting primarily of solitary species, have been recorded from all over the Turkish coastlines (Uysal, 1976; Çınar *et al*., 2006; Okuş *et al*., 2007; Çınar, 2014), in comparison to 45 ascidian species reported from the Levantine basin, along the shores of Israel, Egypt, and the Gulf of Suez (Koukouras *et al*., 1995; Halim & Abdel Messeih, 2016; Reem *et al*., 2017). For updating some of the less studied ascidian species in the Turkish Mediterranean coasts, this study aimed in clarifying the taxonomy, and further apprising the distributions of eight ascidian species by employing COI analyses and major morphological features.

## Materials and Methods

### Sampling and general morphological examinations

Specimens were collected from patchy stony-rocky areas laying on sandy and shallow bottoms (underneath stones, <1 m depth) using razor blades in 6 sites along the Mediterranean coastline of Turkiye, placed, each, in 1,5 mL tube containing 70% ethanol, and kept in room temperature until use (Table S1, Fig. 1). Other colonial fragments were kept in 4% formaldehyde solution at room temperature for morphological analyses. Specimens of all Styelidae species (genera *Botryllus, Botrylloides* and *Symplegma*) were cultured on microscope slides in the Institute of Marine Sciences-Middle East Technical University (IMS-METU) aquaculture room according to Karahan *et al*., (2022) and used for morphological examination. Morphological examinations were carried out for major morphological features (zooid distributions, colonial structures, colors, spicule shape, oral tentacles, blastogenic life cycles), first by the naked eye, followed by stereo and light microscopes (Olympus SZX16 - UC30 camera; Olympus CX43- ToupTek camera).

**Figure 1.**
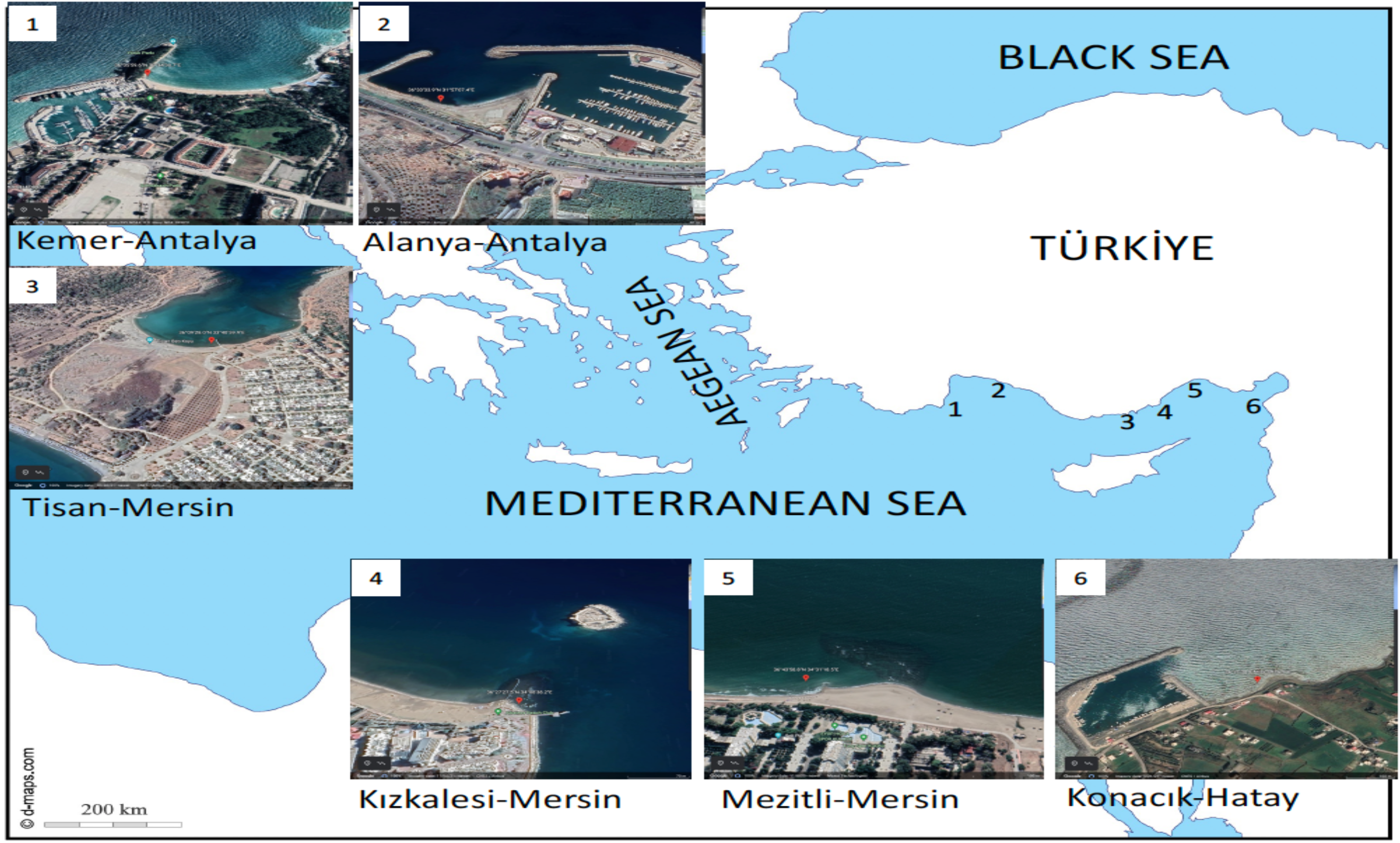
Map and Google Earth captures of the sampling sites. Credits; d-maps.com.

### DNA extraction, Polymerase Chain Reactions (PCR), and data analysis

Total DNA was extracted from colony fragments according to a modified phenol-chloroform method (Karahan *et al*., 2022). Isolated DNA was quantified using a Nanodrop spectrophotometer and diluted, when needed, to 20 ng/μl. PCR were performed in 50 μL total volume with 0.5 μM forward and reverse primers and around 10-20 ng/μl of DNA in a ready-to-use PCR Master Mix (Thermo Scientific) on the mitochondrial cytochrome oxidase subunit I (COI) gene, using Reem et al. (2017) primers (F2-‘AMWAATCATAAAGATATTRGWAC’-3 and R2-‘AARAARGAMGTRTTRAAATTHCGATC’-3). The PCR products were purified and sequenced for forward and reverse directions by Macrogen Inc. (Seoul, South Korea). Detailed data for voucher specimens’ DNA (held in the IMS-METU genetic laboratory) were uploaded to the Barcode of Life Data System (BOLD, http://www.boldsystems.org; Table S2).

BLAST analysis was performed using the GenBank (http://blast.ncbi.nlm.nih.gov/Blast.cgi) and BOLD engines. Sequences were translated and aligned via MAFFTv7 (Katoh *et al*., 2018) and trimmed using Jalview (v 2.11.1.7, Waterhouse *et al*., 2009). The best model for the MrBayes (Ronquist *et al*., 2012) was chosen via PhyML-SMS v3 software (SMS: Smart Model Selection in PhyML; Lefort *et al*., 2017). MrBayes were run according to the GTR+R model for 10.800.000 combined states (two independent runs), resulting in a high effective sample size value (ESS=1829, >100). In total, 45002 trees were sampled after discharging a burn-in fraction of 25% and verifying for LnL stationarity. As convergence diagnostic, we confirmed an average standard deviation of split frequencies below 0.01 (=0.0085), and PSRFs (Potential Scale Reduction Factors) close to 1.0 (Ronquist *et al*., 2012). Final trees were visualized with FigTree v.1.4.4 (Rambaut, 2018, http://tree.bio.ed.ac.uk/software/figtree). MEGA11 (Tamura *et al*., 2021) was used to calculate the Kimura 2-P distance model with 1000 bootstrap and Gamma Distributed rates (Kimura, 1980).

### Database sequences

On total 123 ascidian sequences were mined from NCBI in August 2022 to use species delimitation analyses. Sequences were chosen following the criteria of: having the same genus name, given a voucher record, approved by a taxonomist, and having at least 500 bp length. Yet, few exceptions with sequence lengths between 450 to 500 bp were included in the analyses (like *Botryllus tyreus*-DQ365851-455bp). The IDs of database sequences were given on Fig. 2.

**Figure 2.**
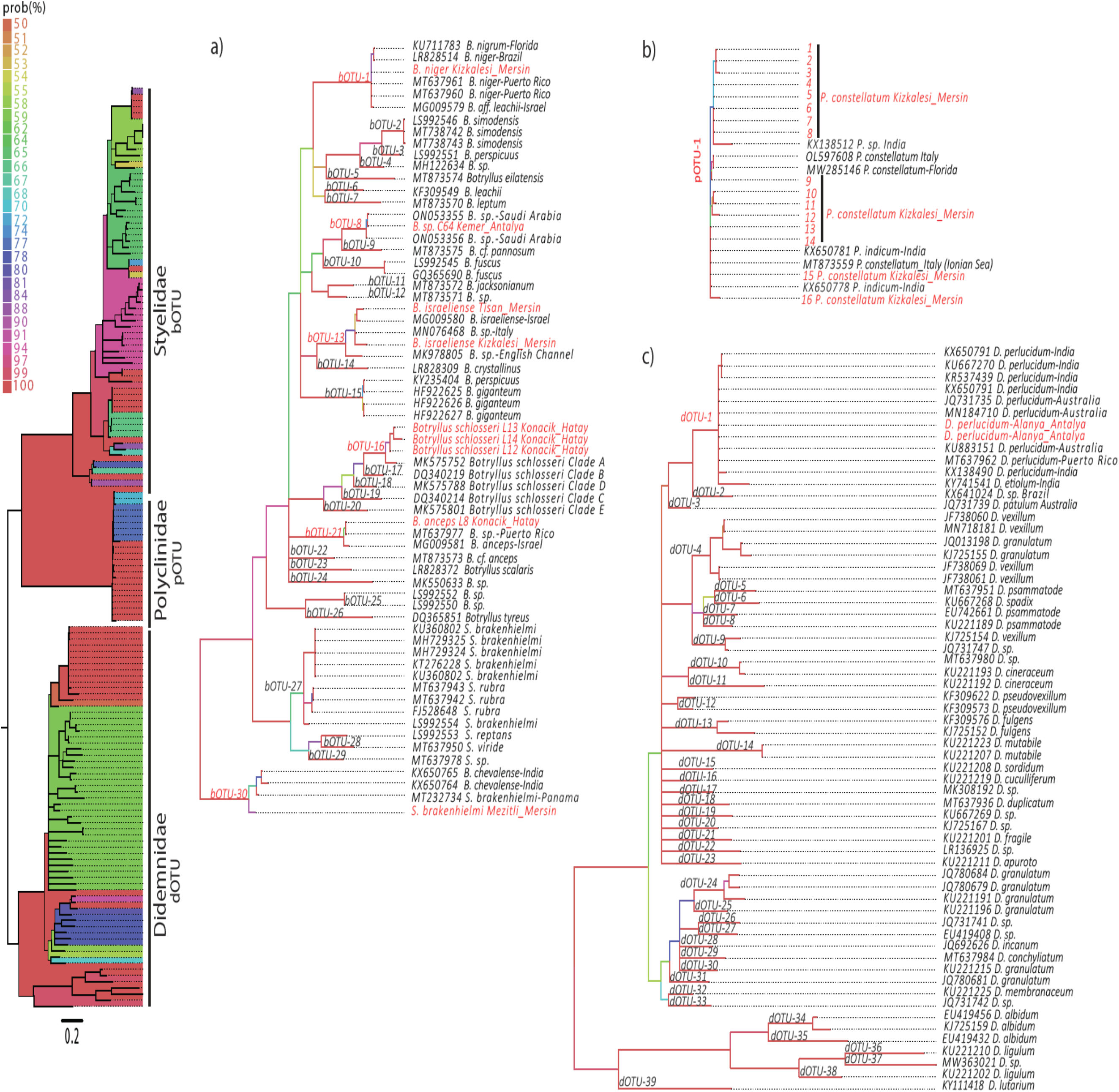
Bayesian majority rule consensus tree reconstructed from the ~ 600 bp COI sequence alignment. a) Styelidae specimens; b) *Polyclinum* specimens; c) Didemnidae specimens. The distance scale is given under the bootstrap probability colored tree, the NCBI IDs are provided next to the species name. OTU numbers are added on the line, according to common results of ASAP and bPTP analyses. Colors of Prob (%) scale, the main tree and a), b) and c) threes’ branches represent probability values. The red colored letters are indicating the present study samples and OTUs. Abbreviations; B.: *Botrylloides*, bOTU: Styelidae family OTUs, pOTU: Polyclinidae family OTUs, dOTU: Didemnidae family OTUs.

### Species delimitation analyses

Analyses were carried out using the Automatic Barcode Gap Discovery method (ASAP; Puillandre *et al*., 2021), a sequence similarity clustering method, and the Poisson Tree Processes (PTP; Zhang *et al*., 2013), a tree-based coalescence method. The hypothetical species identified by these methods were assigned as Operational Taxonomic Units (OTUs). ASAP analyses were performed on the web-based interface (https://bioinfo.mnhn.fr/abi/public/asap/asapweb.html; accessed date: August 2022). Two metric options provided by ASAP were used for the pairwise distance calculations; Jukes-Cantor (JC69; Jukes and Cantor, 1969) and Kimura 2 parameter (K80; Kimura, 1980). PTP analyses were performed using the Bayesian implementation (bPTP), available on the web-based interface (http://species.h-its.org/ptp/, access date: August 2022). MrBayes output (tree) were used in the analyses by performing the default parameter values.

## Results

In total, 27 ascidian sequences from Turkiye were delineated, together with 123 NCBI-mined sequences. According to the common results of ASAP and PTP analyses, in total, 30 OTUs were defined for the genera *Botryllus, Botrylloides* and *Symplegma* (bOTU), 1 OTU for *Polyclinum* (pOTU) and 39 OTUs for *Didemnum* samples (dOTU, Fig. 2 a-c). Present study samples were located in the OTUs of 7 known species, and one sample was positioned with two Saudi Arabia samples as a new *Botrylloides* species (Fig. 2 a,b,c). The morphological and genetic analysis results of each species are detailed below and the ASAP, PTP and LnL scores are found in Fig. S1-S3. Whereas there were common results for most of the OTUs, ASAP assigned less OTU than the PTP in some groups. According to PTP analysis, almost all the *D. perlucidum* samples were assigned to different OTUs, same results were recorded for the *B. schlosseri* samples. The latest OTUs were decided according to all the analysis common results.

### *Botrylloides niger/aff. leachii* (Ascidiacea: Stolidobranchia: Styelidae: Botrylloides)

In total, 221 colony fragments were collected from 6 sampling sites (Table S1), and the sequence of the common haplotype (H1) was used for species delimitation analysis (Temiz *et al*., 2022). Colonies revealed a wide range of color morphs, from a very light color (creamy) to total black (Table S2). The zooids were arranged in ‘leachii type’ systems (Brunetti, 2009), each ca. 1.8 mm long with globular stomachs and same-plane common cloacal opening, where brachial sacs were occasionally seen through it (Fig. 3a_1_-a_4_). Two long (numbers 2-6), six middle (numbers 1, 3, 4, 5, 7 and 8), and many short tentacles containing numerous blood cells fringed the oral siphons (Fig. 3a_4_). The present study samples were assigned in bOTU-1 together with *B. niger/nigrum and B. aff. leachii*, 19% distance from the closest OTUs (bOTU-6 *B. leachii*, Fig. 2a, Table S3).

**Figure 3.**
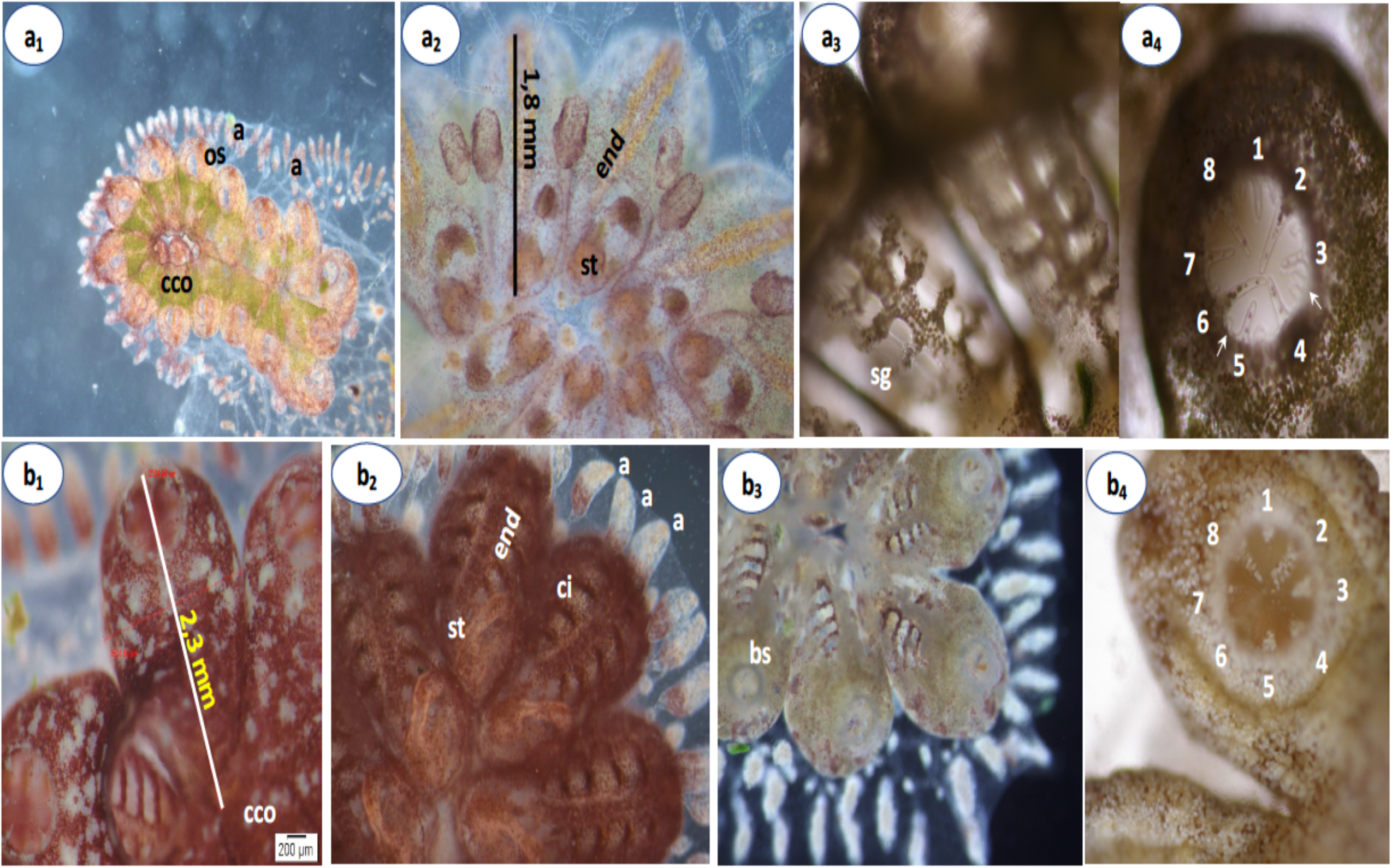
a_1_-a_4_) *Botrylloides niger/aff. leachii*, a_1_) Dorsal and, a_2_) Ventral view of yellow-green color morph of the colony, a_3_) View of brachial sacs through the common cloacal opening, a_4_) Oral siphon and tentacles; 2 long (2 and 6) and 6 middle lengths are numbered, and other short and random tentacles are indicated by arrows. b_1_-b_4_) *Botrylloides israeliense* b_1_) Dorsal view of a zooid, b_2_) Ventral view of the zooids, b_3_) Dorsal view of zooids with brachial sacs (white color morph), b_4_) Numbered oral tentacles (4 long and 4 medium lengths). Abbreviations: end; endostyle, ci: cell island, st; stomach, cco; common cloacal opening, a: ampullas, os: oral siphon

### Botrylloides israeliense

In total, 36 colony fragments with light to dark color morphs (Table S2) were collected from only Mersin sites (Mezitli, Tisan, and Kızkalesi, Table S1), and the sequences of the one common (Kızkalesi sample) and one unique haplotype (Tisan sample) were used for species delimitation analysis. Zooids’ lengths were ca. 2.3 mm, all arranged in a ‘leachii type’ system (Fig. 3b_1_). The stomach had a globular shape continuing after a long esophagus (Fig. 3b_2_). The zooids were positioned vertically to ampullas (Fig. 3b_1_, b_3_). Four long (numbers 2, 4, 6, and 8) and four medium tentacles (numbers 1, 3, 5, and 7) fringed the oral siphons (Fig. 3b_4_).

The sequences of two haplotype (Tisan and Kızkalesi) were used for species delimitation analysis, then were assigned to bOTU-13 with *Botrylloides israeliense* sample from the Mediterranean coast of Israel (NCBI no:MG009580; Reem *et al*., 2017) and two *Botrylloides sp*. samples (MN076468, MK978805, Viard *et al*., 2019) from the English Channel and the Mediterranean Sea, respectively (Fig. 2a). The intra-bOTU-13 distances were 2 to 8%, and the closest OTUs distanced 16 to 19% (bOTU-12, *Botrylloides sp*.) (Fig. 2a, Table S4).

### Botryllus schlosseri

Four colonies collected under two rocks from the Hatay-Konacık region (Table S1), revealed three haplotypes (L12, L13 and L14; Fig. 2a). Haplotype L12 was further cultured in the IMS-METU mariculture system (38-40 ppt salinity and 24-26 °C), subcloned and colonial ramets survived under *in situ* conditions around three years. The colony’s zooids lengths were ca. 1.2 mm revealing a brown color morph with yellow stripes (Fig. 4a_1_, 4a_2_). In time the pigmentation increased, and the blastogenic cycles became irregular.

**Figure 4.**
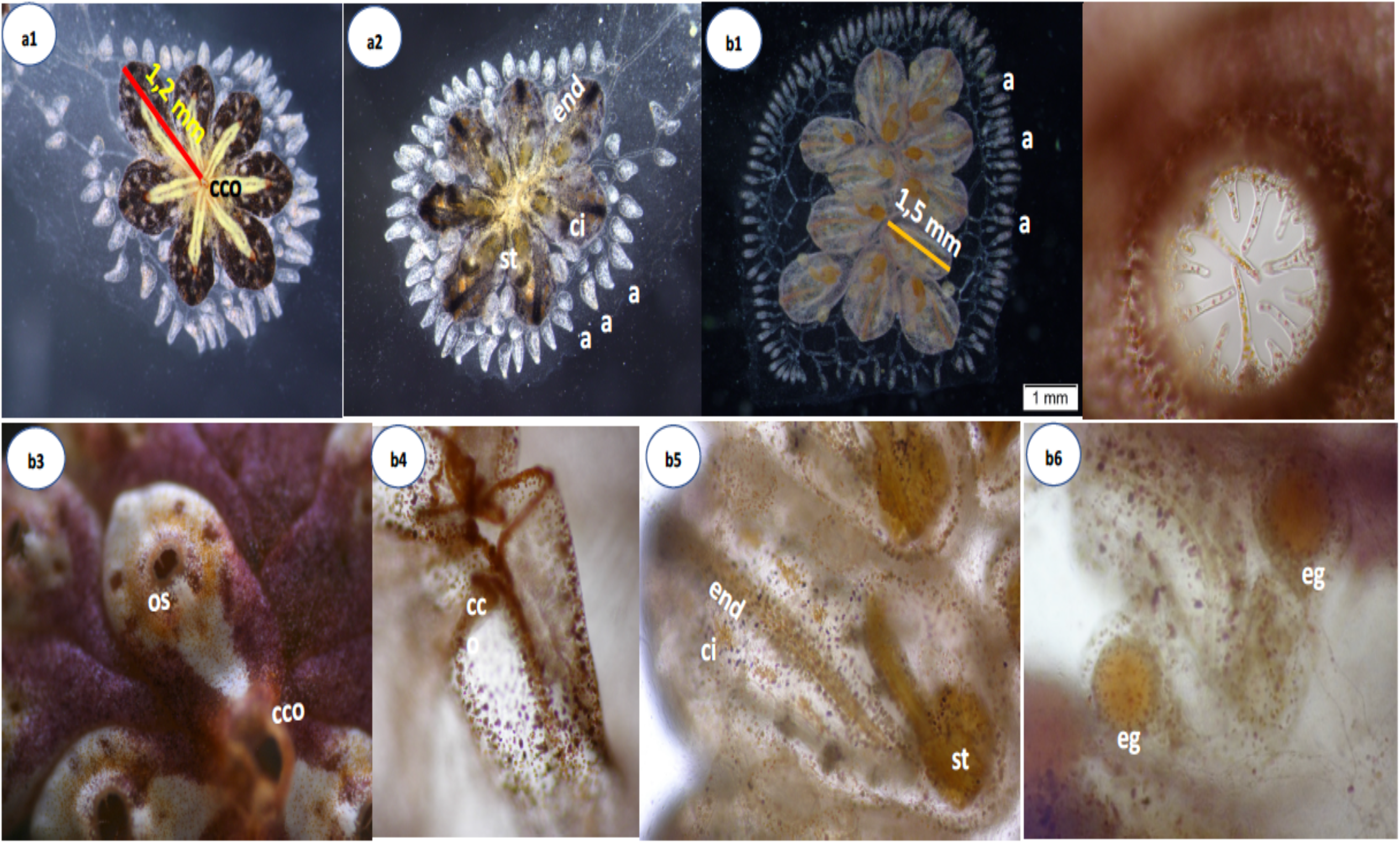
a_1_-a_2_) *Botryllus schlosseri* a_1_) Dorsal and a_2_) Ventral view of a cultured colony. b_1_-b_6_) *Botrylloides anceps*: b_1_) Ventral view of a pink color morph b_2_) Oral siphon and tentacles (6 long, 6 short and few very short), b_3_) Dorsal view of a purple color morph zooid b_4_) Common cloacal opening (cco), b_5_) Single zooid ventral view, b_6_) Primary bud with eggs. Abbreviations: end; endostyle, ci: cell island, st; stomach, eg; egg, cco; common cloacal opening, a: ampullas, bv; blood vessels, vb; vascular bud, os: oral siphon, co; cloacal opening.

All three Turkish haplotypes were assigned into bOTU-16 as Clade A colonies together with a database sample (MK575752). Other *B. schlosseri* samples mined from GeneBank were assigned to four different OTUs; bOTU-17, −18, −19, and −20 (Table S5). The intra-bOTU-16 distance varied between 2 and 5%, and the highest distance was noted between our L14 sample and Clade E (bOTU-20) at 19.5 % (Fig. 2a, Table S5).

### Botrylloides anceps

In total 27 colonial samples were collected from Hatay-Konacık, Mezitli-Mersin, and Alanya-Antalya regions (Fig. 1, Table S1) and the sequence of the one common haplotype was used for species delimitation analysis. Zooids (between 1.5-2 mm long, Fig. 4 and S4) were organized in ‘leachii type’ systems and color morphs vary from very light pink to dark brown, and creamy to purple (Fig. 4b_1_, b_3_). Two long, four middle, and six short size tentacles fringed the oral siphons (Fig. 4b_2_). The common cloacal opening is quite large, serving around 14 zooids (Fig. 4b_4_). Stomachs with globular shapes (Fig. 4b_5_). Laboratory follow-up observations revealed sexual reproduction under 38-40 ppt salinity and 26-28 °C, where eggs were recorded in zooids and primary buds, and testes on zooids (Fig. 4b_6_). While most progenies died at the oozooid stage, some survived 2-3 years under the IMS-METU mariculture conditions (16 to 32 °C), while young colonies grew, without signs for morphological hibernation (sensu Hyams *et al*., 2017, 2022), and 2-3 years old colonies showed exaggerated, yet condensed ampullae and irregular blastogenic cycles (all resembling onset of hibernation), a few months before their death (Fig. S5).

All 27 *B. anceps* samples belong to a single haplotype, and the Hatay region sample sequence was assigned to bOTU-21 together with the Israeli *B. anceps* (MG009581, Reem *et al*., 2017) and *Botrylloides sp*. from Puerto Rico (MT637977; Streit *et al*., 2021), differentiating with less than 0.01%. *Botrylloides cf. anceps* (MT873573, Salonna *et al*., 2021) was assigned to another OTU (bOTU-22), ca. 18 % distance from bOTU-21 (Fig. 2a, Table S6).

### Botrylloides sp

A single colony (brownish with a yellow stripe around the oral siphon and dorsal site) of ‘leachii type’ systems (Fig. 5a_1_) was collected from the Antalya-Kemer region (Table S1). A colony fragment (25 zooids) was cultured for one month, revealing 1.7 mm long zooids (Fig. 5a_2_, a_3_) and blastogenic cycles of five days long under 26°C and 38-40 ppt salinity (Fig. S6).

**Figure 5.**
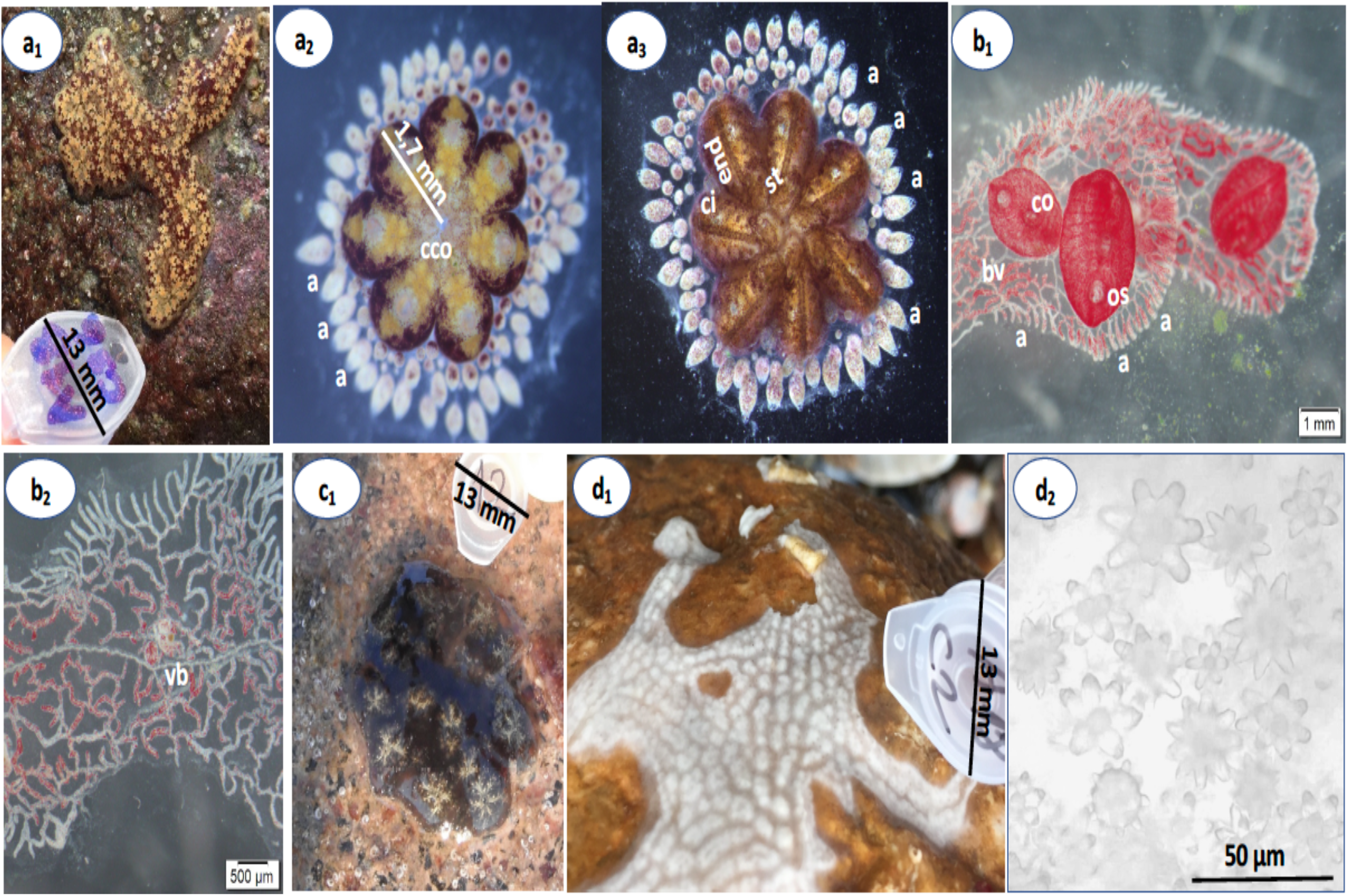
a_1_-a_3_) *Botrylloides sp*. a_1_) field view a_2_) cultured colony’s dorsal and a_3_) ventral view. b_1_-b_2_) *Symplegma brakenhielmi*; b_1_) Dorsal and ventral views of a colony, b_2_) Blood vessels and palleal budding, c_1_) *Polyclinum constellatum* dorsal view, d_1_-d_2_) *Didemnum perlucidum*; d_1_) Dorsal view of a colony, d_2_) the spicules of the colony. Abbreviations: end; endostyle, ci: cell island, st; stomach, eg; egg, cco; common cloacal opening, a: ampullas, bv; blood vessels, vb; vascular bud, os: oral siphon, co; cloacal opening.

This colony was assigned to bOTU-8 together with two database samples from Saudi Arabia, Yanbu (*Botrylloides sp*.; ON053355-ON053356). The closest OTU (bOTU-9, *Botrylloides cf. pannosum*, MT873575; Bari, Italy; Salonna *et al*., 2021) is of a 14% distance (Fig 2a, Table S7).

### *Symplegma brakenhielmi* (Ascidiacea; Stolidobranchia; Styelidae; Symplegma)

A single red colony was collected from the Mersin-Mezitli region under a rock (Fig. 1, Table S1). A ramet cultured in the aquaculture room (38-40 ppt, 27-28 °C; Fig. 5b_1_) revealed transparent tunic, with dispersed zooids arranged with no apparent systems. Both palleal and vascular budding were recorded, with the latter more prominent, 1 cm long ampullae and 1-3 cm long adult zooids (Fig. 5b_1_-b_2_).

This specimen was assigned to bOTU-30 as *S. brakenhielmi* together with a Panama sample (MT232734) at a 3% distance (Table S8, Fig. 2a) and two *Botrylloides chevalense* samples (KX650765 and KX650764) from India, yet with a 4-5% distance (Table S8, Fig. 2a). Other *S. brakenhielmi* samples, four Indian (KU360802, MH729325, MH729324, KT276228) and one from Italy (Bari; LS992554) samples were assigned to a different OTU (bOTU-27), together with *S. rubra* samples, ca. 28% distance to bOTU-30 (Fig. 2a, Table S8).

### Polyclinum constellatum (Ascidiacea: Aplousobranchia: Polyclinidae: Polyclinum)

Sixteen colonies (light brown, reddish, and gray with a beige system of zooids) were collected only Kızkalesi site in 2014 (Fig. 1, Fig. 5c_1_).

These samples belonged to two haplotypes and were assigned into a single OTU (pOTU-1) together with three database samples of *P. constellatum* from Italy and Florida (OL597608; MT873559; MW285146), and with *P. indicum* and *Polyclinum sp*. (KX650781; KX650778; KX138512) from India, with 6% intra- pOTU-1 distance (Table S9, Fig. 2b).

### *Didemnum perlucidum* (Ascidiacea: Aplousobranchia: Didemnidae:*Didemnum*)

This species was commonly observed on hard substrates of all sites (white, orange, light and dark creamy, and brownish color morphs). Colonies were less than 1mm thickness (Fig. 1; Table S1; Fig. 5d_1_, Table S2) and with star-shaped spicules (Fig. 5d_2_).

COI sequences from two colonies (from Alanya) were used for species delimitation analysis, together with the GeneBank data. In total, 39 OTUs were assigned for all *Didemnum* species (Fig. 2c). According to ASAP score present study samples were clustered under the dOTU-1 together with *D. perlucidum* (Fig2c, Table S10) and *D. etiolum* (KY741541) samples from India, Australia and Puerto Rico with between the 0% and 5.7% distances (Table S10).

## Discussion

This study reveals the existence and spatial distribution of eight colonial ascidian species from the shallow Turkish North Eastern Mediterranean coastlines. With the current sampling effort (focused on the very shallow waters), there is a high possibility that not all colonial species were sampled due to seasonality or deeper ascidian habitats. In total, five botryllid ascidians and one *Symplegma*, one *Polyclinum*, and one *Didemnum* species were recorded from six sites between Hatay and Antalya coastlines. Distance matrix, Bayesian tree, and PTPs and ASAP analyses scores were used for species assignment. Whereas more OTUs were recorded in PTP than the ASAP, common OTUs were chosen and differences were connected to data set size that was also experienced in the previous study (Goulpeau *et al*., 2022).

*Botrylloides niger/aff. leachii* represents one of the emerged ambiguities in the taxonomic assignment and species delineation in botryllid ascidians (Reem *et al*., 2018). The first record of *Botrylloides niger* Herdman, 1886 for the Mediterranean Sea was given by Peres (1958), and its wide distribution was reported from the Atlantic, Pacific, and Indian Ocean, Mediterranean Sea, and the Red Sea (https://www.marinespecies.org/aphia.php?p=taxdetails&id=252289). Based on mtCOI sequences, recent studies (Goutman *et al*., 2020; Temiz *et al*., 2022) reassigned the dwarf form of *Botrylloides leachii* from the Levant (Brunetti, 2009) as *Botrylloides nigrum*. A study (Reem *et al*., 2018) employing of three additional markers (18S, 28S, H3) contradicted literature suggestions, revealing minute distances between *Botrylloides leachii* and the ‘dwarf *Botrylloides leachii*’. As this issue has not yet been solved, we assigned this species here as *Botrylloides niger/aff. leachi*. Colonies of this species were collected from all the sampling sites, representing diverse color morphotypes previously recorded in the Mediterranean Sea (Rinkevich *et al*., 1993; Brunetti, 2009), the Pacific and Atlantic oceans (Goodbody, 2000, 2003; Monniot & Monniot 1994).

*Botrylloides israeliense* is a newly identified colonial ascidian (Brunetti, 2009) from the Mediterranean coast of Israel (Bay of Àkko and Mikhmoret) that was already studied in the past for some ecological and biological characteristics (Rinkevich *et al*., 1993,1994).The genetic distances between the Turkish and the Israeli samples were between 3 to 5%, a value that is above the general eukaryotic species threshold (<2%, Hebert *et al*., 2004) but still lower than some other cases (~20% Resch et al., 2014; Reem *et al*., 2022).

*B. schlosseri* is a highly invasive colonial ascidian distributed worldwide primarily in temperate shallow waters. Whereas the presence of the species on the Turkish coasts was reported before (Çınar, 2014; Kayış, 2011), no DNA sequence data was provided. So far, five clades have been assigned (A-E) worldwide (Reem *et al*., 2022), and the present study samples clustered into the most common clade (A) with up to a 5.2% intra-clade distance as compared to the worldwide within clade A distance up to 6.3%. Further, according to ASAP and pPTP results clades B, C, D, and E were assigned as different species with min.11.2% and max. 17.6% distance scores, a result that casts a query about the validity of current delimitation analyses.

*Botrylloides anceps* is also believed to be a NIS in the Mediterranean Sea. The first record was given from the Israeli coasts by Brunetti (2009). We recorded the species for the first time in 2018 at Hatay, Mezitli, and Alanya sites. A previous visit to the sites had been done in 2012 and no single *B. anceps* individuals were recorded, suggesting a recent introduction by the low (0.7%) distance between the Israeli and the Turkish samples. One important morphological characteristic of this study is the variable sizes assigned to *B. anceps* zooids, a morphometric quality that is changed according to the colony size, adding difficulties to this simple taxonomic characteristic for ascidian species identifications.

Colonies of presumably a new species *Botrylloides sp*. were recorded from only Antalya-Kemer sampling locations. This species matched 99% the NCBI samples uploaded from Saudi Arabia and the distance between the closest *Botrylloides* relatives was assigned at 14% (*Botrylloides cf. pannosum*; Australia). While this new botryllid ascidian species may have originated from the Red Sea, it has not been recorded from the Mediterranean coasts of Israel and the east sites of the Turkiye (Hatay and Mersin), thus making its origin fuzzy.

The first record of *Symplegma brakenhielmi* in the Mediterranean Sea was from the Lebanese coast, then from the Turkish Levantine coast (Bitar & Kouli-Bitar, 2001; Çınar *et al*., 2006), based on only morphological characteristics. Other records included the central part of the Mediterranean Sea (Ulman *et al*., 2017; Mastrototaro *et al*., 2019). Our study on the Northeastern Mediterranean Sea reveals, for the first time, a species delimitation analysis that further indicated a possible misidentification and sequencing errors in the database, as the Turkish colonies were clustered with Panamian *S. brakenhielmi* and Indian *B. chevalense* samples, and were distinct from the Mediterranean (Bari, Italy) *S. brakenhielmi* cluster. Moreover, the present study sample is located as the most ancestral lineage on the tree with 100% bootstrap support which may indicate polyphyly for the genus. On the other hand, like previous study (Mastrototaro *et al*., 2019) *S. brakenhielmi* and *S. rubra* clustered together under the same OTU (bOTU-27), which is refer to the necessity of further morphological and genetic analysis for the whole genus.

With its wide range distribution (Western Atlantic, Western Indian Ocean, Indo-Pacific, Mediterranean and Aegean), *Polyclinum constellatum* has been considered a cryptogenic species (Dias *et al*., 2013; Halim and Messeih, 2016; Aydın-Önen, 2018; Montesanto *et al*., 2022). Recording another name (*P. indicum)* under the *P. constellatum* OTU (pOTU1) was referred to as an error or synonymy (Montesanto *et al*., 2022). Present study samples were highly matched (99-100%) with database specimens uploaded from various locations all over the world (Italy, Florida, India, and the Ionian Sea), a result indicating low intraspecific distance in a widely spread species and also monophyly.

*Didemnum perlucidum* is also a widespread species (Atlantic, Pacific, Indian Oceans, and Mediterranean Sea), and its native range is unknown (Lambert, 2002; Muñoz *et al*., 2015; Novak & Shenkar, 2020). Here we provide the first barcode data for the Northeastern Mediterranean Sea populations and a high-resolution phylogenetic tree, revealing genetic distances (0 to 6%) between the widely disturbed populations (Turkiye, India, Australia, Porto-Rico). Further, whereas the Turkish samples are clustered in a single OTU together with other *Didemnum perlucidum* samples, other valid species were assigned in the same OTU (like dOTU4, *Didemnum vexillum*, and *Didemnum granulatum*), while samples of other inclusive species i (e.g., *Didemnum psammatode*) were further clustered under different OTUs (like dOTU 5, 7, and 8). All above highlighted the needs for taxonomic revision.

In conclusion, we provide here the descriptions for eight colonial ascidians from the Turkish North Eastern Mediterranean Sea shallow waters, including barcode data and species delimitation, supported with major morphological features of studied species. Besides high variation in morphotypes (*B. niger/aff. leachii* and *B. israeliense*), high (*B. schlosseri*) and low (*Didemnum perlucidum* and *P. constellatum*) genetic diversities were also recorded. A possible new *Botrylloides* species (*Botrylloides sp*.) from the Antalya region was revealed. It is also understood that the zooid length cannot always be used as an informative taxonomic feature for *B. anceps*. From a phylogenetic perspective, *Polyclinum* members were found closer to the Styelidae members than Didemnidae and a possible polyphyly was recorded for the *Symplegma* genus and Didemnidae family.

## Supporting information

Karahan_Supplementary data

## Acknowledgments

Thanks to the municipalities of Erdemli and Manavgat for their help during the sampling. We thank the Institute of Marine Sciences-METU for providing transportation during our field trips. We also thank Kenan Murat Karahan, Fatima Nur Oğul and Begüm Ece Tohumcu for their help during the sampling. This study was supported by grants from the YÖP-701-2018-2666, Middle East Technical University support program, and the research was executed by using “IMS-METU, DEKOSIM laboratory (BAP-08-11-DPT2012K120880) facilities.

## Supplementary Materials

Table S1. Sampling sites and date details

Table S2. General information of BoLD uploaded and other specimens

Table S3. Kimura-2 Parameter distance for bOTU-1 and −7.

Table S4. Kimura-2 Parameter distance for bOTU-13 and −12.

Table S5. Kimura-2 Parameter distance for bOTU-16, −17, −18, −19 and −20.

Table S6. Kimura-2 Parameter distance for bOTU-21 and −22

Table S7. Kimura-2 Parameter distance for bOTU-8 and −9.

Table S8. Kimura-2 Parameter distance for bOTU-27 and −30.

Table S9. Kimura-2 Parameter distance for pOTU-1

Table S10. Kimura-2 Parameter distance for dOTU-1

Fig. S1. ASAP score; Colors represent different OTUs. Number in line of the OTUs presents the total samples assigned to the same OTU. The first number line above the OTUs’ columns presents the total OTU numbers, values at the second line indicate ASAP-scores (the lowest the score the better is the partition, Puillandre et al. 2021).

Fig. S2. PTP score; Blue lines present different OTUs, red lines present same OTUs

Fig. S3. LnL value of MrBayes analysis.

Fig. S4. When the zooid size of the brown morphotypes of *B. anceps* reached up to 2 mm.

Fig. S5. Large, retracted and condensed ampullas of *Botrylloides anceps*

Fig. S6. Blastogenic cycle of *Botrylloides sp*. Dorsal and ventral view of the Blastogenic stages; a-b) stage-D (Jun 07, 2018), c-d) stage-A (Jun 08, 2018), e-f) stage-B (Jun 09, 2018), g) Dorsal-stage C (Jun 10, 2018), h) Dorsal – stage D (Jun 11, 2018). Abbreviations: end; endostyle, st; stomach, cco; common cloacal opening, sg: stigmata, a: ampullas.

## Supplementary Materials

Ten supplementary tables and six supplementary figures are provided as supporting information.

## Author Contributions

Conceptualization, A.K.; Data curation, A.K., B.T., E.Ö., J.D., and B.R.; Methodology, A.K., B.T., and E.Ö.; Writing-original draft, A.K., B.T., E.Ö., JD., and BR. All authors have read and agreed to the published version of the manuscript.

## Data Availability Statement

Sequences, trace files, image files, and the primers information for each COI haplotype were uploaded to the Barcode of Life Data System (Ratnasingham & Hebert, 2007). All the data will be public upon acceptance.

## References

Aydın-Önen, S.A., 2018. Distribution of ascidians with a new record of the non-indigenous species *Polyclinum constellatum* savigny, 1816 from the Aegean coast of Turkey. Turkish Journal of Fisheries and Aquatic Sciences, 18 (9), 1077–1089. https://doi.org/10.4194/1303-2712-v18_9_07

Berrill, N.J., 1950. The Tunicata with an account of the British species. Ray Society, London, 354 pp.

Bitar, G., Kouli-Bitar, S., 2001. Nouvelles données sur la faune et la flore benthiques de la cote Libanaise. Migration Lessepsienne [New data on benthic fauna and flora on the coast Lebanese. Lessepsian migration]. Thalassia Salentina, 25, 71–74.

Brunetti, R., 2009. Botryllid species (Tunicata, Ascidiacea) from the Mediterranean coast of Israel, with some considerations on the systematics of Botryllinae. Zootaxa, 2289 (1). https://doi.org/10.11646/zootaxa.2289.1.2

Brunetti, R., Manni, L., Mastrototaro, F., Gissi, C., Gasparini, F., 2017. Fixation, description and DNA barcode of a neotype for *Botryllus schlosseri* (Pallas, 1766) (Tunicata, Ascidiacea). Zootaxa, 4353 (1), 29–50.

Burnet, F.M., 1971. “Self-recognition” in colonial marine forms and flowering plants in relation to the evolution of immunity. Nature, 232 (5308), 230–235. https://doi.org/10.1038/232230a0

Çınar, M.E., Bilecenoglu, M., Öztürk, B., Can, A., 2006. New records of alien species on the Levantine coast of Turkey. Aquatic Invasions, 1 (2), 84–90. https://doi.org/10.3391/ai.2006.1.2.6

Çınar, M.E., 2014. Checklist of the phyla platyhelminthes, Xenacoelomorpha, Nematoda, Acanthocephala, Myxozoa, Tardigrada, Cephalorhyncha, Nemertea, Echiura, Brachiopoda, Phoronida, Chaetognatha, and chordata (Tunicata, Cephalochordata, and hemichordata) from the coasts of Turkey. Turkish Journal of Zoology, 38 (6), 698–722. https://doi.org/10.3906/zoo-1405-70

Denoeud, F., Henriet, S., Mungpakdee, S., Aury, J.M., da Silva, C. et al., 2010. Plasticity of animal genome architecture unmasked by rapid evolution of a pelagic tunicate. Science, 330 (6009), 1381–1385. https://doi.org/10.1126/science.1194167

Dias, G.M., Rocha, R.M., Lotufo, T.M.C., Kremer, L.P., 2013. Fifty years of ascidian biodiversity research in São Sebastião, Brazil. Journal of the Marine Biological Association of the United Kingdom, 93 (1), 273–282. https://doi.org/10.1017/S002531541200063X

Galil, B.S., Marchini, A., Occhipinti-Ambrogi, A., 2018. East is east and West is west? Management of marine bioinvasions in the Mediterranean Sea. Estuarine, Coastal and Shelf Science, 201, 7–16. https://doi.org/10.1016/j.ecss.2015.12.021

Goodbody, I., 2000. Diversity and distribution of ascidians (Tunicata) in the Pelican Cays, Belize. Atoll Research Bulletin, 480, 301–326.

Goodbody, I., 2003. The ascidian fauna of Port Royal, Jamaica I. Harbor and mangrove dwelling species. Bulletin of Marine Science, 73 (2), 457–476.

Goulpeau, A., Penel, B., Maggia, M.E., Marchán, D.F., Steinke, D. et al., T., 2022. OTU Delimitation with Earthworm DNA Barcodes: A Comparison of Methods. Diversity, 14, 866. https://doi.org/10.3390/d14100866

Goutman, S.A., Boss, J., Guo, K., Alakwaa, F.M., Patterson, A. et al., 2020. Untargeted metabolomics yields insight into ALS disease mechanisms. Journal of Neurology, Neurosurgery, and Psychiatry, 91 (12), 1329–1338. doi: 10.1136/jnnp-2020-323611.

Halim, Y., Abdel Messeih, M., 2016. Aliens in Egyptian waters. A checklist of ascidians of the Suez Canal and the adjacent Mediterranean waters. Egyptian Journal of Aquatic Research, 42 (4), 449–457. https://doi.org/10.1016/j.ejar.2016.08.004

Hebert, P.D.N., Stoeckle, M.Y., Zemlak, T.S., Francis, C.M., 2004. Identification of birds through DNA Barcodes. PLoS Biology, 2 (10), e312. https://doi.org/10.1371/journal.pbio.0020312

Hebert, P.D., Cywinska, A., Ball, S.L., deWaard, J.R., 2003. Biological identifications through DNA barcodes. Proceedings of Royal Society B: Biological Science, 270, 313–21

Hyams, Y., Paz, G., Rabinowitz, C., Rinkevich B., 2017. Insights into the unique torpor of *Botrylloides leachi*, a colonial urochordate. Developmental Biology, 428 (1), 101–117.

Hyams, Y., Panov, Y., Rosner, A., Brodski, L., Rinkevich, Y. et al., 2022. Transcriptome landscapes that signify *Botrylloides leachi* (Ascidiacea) torpor states. Developmental Biology, 490, 22–36.

Jukes, T.H., Cantor, C.R., 1969. Evolution of protein molecules. Academic Press. New York and London, 412 pp.

Karahan, A., Öztürk, E., Temiz, B., Blanchoud, S., 2022. Studying tunicata WBR using *Botrylloides anceps*. p. 311–332. In: Whole-Body Regeneration. Methods in Molecular Biology. Vol. 2450. Blanchoud, S., Galliot, B. (Eds). Humana Press, New York. doi: 10.1007/978-1-0716-2172-1_16

Karahan, A., Douek, J., Paz, G., Rinkevich, B., 2016. Population genetics features for persistent, but transient, *Botryllus schlosseri* (Urochordata) congregations in a central Californian marina. Molecular Phylogenetics and Evolution, 101, 19–31. https://doi.org/10.1016/j.ympev.2016.05.005

Katoh, K., Rozewicki, J., Yamada, K.D., 2018. MAFFT online service: multiple sequence alignment, interactive sequence choice and visualization. Briefings in Bioinformatics, 20, 1160–1166

Kayış, Ş., 2011. Ascidian Tunicate, *Botryllus schlosseri* (Pallas, 1766) infestation on seahorse. Bulletin of the European Association of Fish Pathologists, 18 (2), 81–84.

Kimura, M.A., 1980. Simple method for estimating evolutionary rates of base substitutions through comparative studies of nucleotide sequences. Journal of Molecular Evolution, 16, 111–120.

Koukouras, A., Voultsiadou-Koukoura, E., Kevrekidis, T., Vafidis, D., 1995. Ascidian fauna of the Aegean Sea with a check list of the eastern Mediterranean and Black Sea species. Annales de l’Institute Oceanographique, 71 (1), 19–34.

Lambert, G., 2002. Nonindigenous ascidians in tropical waters. Pacific Science, 56 (3), 291–298. https://doi.org/10.1353/psc.2002.0026

Lefort V., Longueville J-E., Gascuel O., 2017. SMS: Smart Model Selection in PhyML. Molecular Biology and Evolution, 34 (9), 2422–2424.

López-Legentil, S., Legentil, M.L., Erwin, P.M., Turon, X., 2015. Harbor networks as introduction gateways: contrasting distribution patterns of native and introduced ascidians. Biological Invasions, 17 (6), 1623–1638. https://doi.org/10.1007/s10530-014-0821-z

Mastrototaro, F., Montesanto, F., Salonna, M., Grieco, F., Trainito, E. et al., 2019. Hitch-hikers of the sea: Concurrent morphological and molecular identification of *Symplegma brakenhielmi* (Tunicata: Ascidiacea) in the western Mediterranean Sea. Mediterranean Marine Science, 20 (1), 197–207. https://doi.org/10.12681/mms.19390

Monniot, C., Monniot, F., 1994. Additions to the inventory of eastern tropical Atlantic ascidians; arrival of cosmopolitan species. Bulletin of Marine Science, 54 (1), 71–93.

Montesanto, F., Chimienti, G., Gissi, C., Mastrototaro, F., 2022. *Polyclinum constellatum* (Tunicata, Ascidiacea), an emerging non-indigenous species of the Mediterranean Sea: integrated taxonomy and the importance of reliable DNA barcode data. Mediterranean Marine Science, 23 (1), 69–83. https://doi.org/10.12681/mms.28311

Muñoz, J., Page, M., McDonald, J.I., Bridgwood, S.D., 2015. Aspects of the growth and reproductive ecology of the introduced ascidian *Didemnum perlucidum* (Monniot, 1983) in Western Australia. Aquatic Invasions, 10 (3), 327–332. https://doi.org/10.3391/ai.2015.10.3.02

Novak, L., Shenkar, N., 2020. Occurrence of *Didemnum perlucidum* Monniot F., 1983 on artificial substrates along the Mediterranean coast of Israel. Mediterranean Marine Science, 21 (2), 386–392. https://doi.org/10.12681/mms.22223

Okuş, E., Altıok, H., Yüksek, A., Yılmaz, N., Yılmaz, A. et al., 2007. Biodiversity in western part of the Fethiye Bay. Black Sea/Mediterranean Environment, 13, 19–34

Peres, J.M., 1958. Ascidies recoltees sur les cotes Mediterraneenaires d’Israel. Bulletin of the Research Council of Israel, 7B, 143–150.

Puillandre, N., Brouillet, S., Achaz, G., 2021. ASAP: assemble species by automatic partitioning. Molecular Ecology Resources, 21 (2), 609–620. https://doi.org/10.1111/1755-0998.13281

Rambaut, A., 2018. FigTree: tree figure drawing tool version 1.4.4. http://tree.bio.ed.ac.uk/software/figtree/.

Ratnasingham, S., Hebert, P.D.N., 2007. BOLD: The barcode of life data system (http://www.barcodinglife.org). Molecular Ecology Notes, 7 (3), 355–364. https://doi.org/10.1111/j.1471-8286.2007.01678.

Reem E., Douek, J., Rinkevich, B., 2017. Ambiguities in the taxonomic assignment and species delineation of botryllid ascidians from the Israeli Mediterranean and other coastlines. Mitochondrial DNA Part A, 29 (7), 1073–1080.

Reem, E., Douek, J., Rinkevich, B. 2018. Ambiguities in the taxonomic assignment and species delineation of botryllid ascidians from the Israeli Mediterranean and other coastlines. Mitochondrial DNA Part A: DNA Mapping, Sequencing, and Analysis, 29(7), 1073–1080. https://doi.org/10.1080/24701394.2017.1404047

Reem, E., Douek, J., Rinkevich, B., 2022. A critical deliberation of the “species complex” status of the globally-spread colonial ascidian *Botryllus schlosseri*. Journal of the Marine Biological Association of the United Kingdom, 101, 1047–1060. https://doi.org/10.1017/S0025315422000029

Resch, M.C., Shrubovych, J., Bartel, D., Szucsich, N.U., Timelthaler, G. et al., 2014. Where taxonomy based on subtle morphological differences is perfectly mirrored by huge genetic distances: DNA barcoding in Protura (Hexapoda). PLoS One, 9 (3), e90653. https://doi.org/10.1371/journal.pone.0090653

Rinkevich, B., Lilker-Levav, T., Goren, M., 1994. Allorecognition/xenorecognition responses in *Botrylloides* (Ascidiacea) subpopulations from the Mediterranean coast of Israel. Journal of Experimental Zoology, 270 (3), 302–313.

Rinkevich, B., Shlemberg, Z., Lilker-levav, T., Goren, M., Fishelson, L., 1993. “Life history characteristics of *Botrylloides* (Tunicata) populations in Akko Bay, Mediterranean Coast of Israel.” Israel Journal of Zoology, 39 (3), 197–212.

Rocha, R.M., da Zanata, T.B., Moreno, T.R., 2012. Keys for the identification of families and genera of Atlantic shallow water ascidians. Biota Neotropica, 12 (1). https://doi.org/10.1590/s1676-06032012000100022

Ronquist, F., Teslenko, M., van der Mark, P., Ayres, D.L., Darling, A. et al., 2012. Mrbayes 3.2: Efficient bayesian phylogenetic inference and model choice across a large model space. Systematic Biology, 61 (3), 539–542. https://doi.org/10.1093/sysbio/sys029

Rubinstein, N.D., Feldstein, T., Shenkar, N., Botero-Castro, F., Griggio, F. et al., 2013. Deep sequencing of mixed total DNA without barcodes allows efficient assembly of highly plastic ascidian mitochondrial genomes. Genome Biology and Evolution, 5 (6), 1185–1199. https://doi.org/10.1093/gbe/evt081

Salonna, M., Gasparini, F., Huchon, D., Montesanto, F., Haddas-Sasson, M. et al., 2021. An elongated COI fragment to discriminate botryllid species and as an improved ascidian DNA barcode. Scientific Reports, 11 (1), 1–19. https://doi.org/10.1038/s41598-021-83127-x

Savigny, J.C., 1816. Mémoires Sur Les Animaux Sans Vertèbres. Paris, 240 pp.

Schlosser, J.A., Ellis, J., 1755. An account of a curious, fleshy, coral-like substance; in a letter to Mr. Peter Collinson, F. R. S. from Dr. John Albert Schlosser, M. D. F. R. S. with some observations on it communicated to Mr. Collinson by Mr. John Ellis, F. R. S. Philosophical Transactions, 49, 449–452.

Spallanzani, L., Chiereghin, S., 1784. Botryllus schlosseri. https://sites.google.com/site/ascidianbiologylab/clients (Accessed 23 December 2022)

Streit, O.T., Lambert, G., Erwin, P.M., López-Legentil, S., 2021. Diversity and abundance of native and non-native ascidians in Puerto Rican harbors and marinas. Marine Pollution Bulletin, 167, 112262. https://doi.org/10.1016/j.marpolbul.2021.112262

Tamura, K., Stecher, G., Kumar, S., 2021. MEGA11: Molecular evolutionary genetics analysis version 11. Molecular Biology and Evolution, 38 (7), 3022–3027.

Temiz, B., Öztürk, E., Blanchoud, S., Karahan, A., 2022. Phylogeographic and morphological analysis of *Botrylloides niger* Herdman, 1886 from the northeastern Mediterranean Sea. BioRxiv, 1. doi: 10.1101/2022.11.30.518487 (Journal revision is completed)

Ulman, A., Ferrario, J., Occhpinti-Ambrogi, A., Arvanitidis, C., Bandi, A. et al., 2017. A massive update of non-indigenous species records in Mediterranean marinas. PeerJ, 5, e3954.

Uysal, A., 1976. Ascidians in Turkish Waters (in Turkish). İ.Ü. Fen Fakültesi Hidrobioloji, Araştırma Enstitüsü Yayınları. 15, 29.

Viard, F., Roby, C., Turon, X., Bouchemousse, S., Bishop, J., 2019. Cryptic diversity and database errors challenge non-indigenous species surveys: An illustration with *Botrylloides spp*. in the English Channel and Mediterranean Sea. Frontiers in Marine Science, 6, 615. https://doi.org/10.3389/fmars.2019.00615

Waterhouse, A.M., Procter, J.B., Martin, D.M.A., Clamp, M., Barton, G.J., 2009. Jalview Version 2-A multiple sequence alignment editor and analysis workbench. Bioinformatics, 25 (9), 1189–1191. https://doi.org/10.1093/bioinformatics/btp033

Zenetos, A., Çinar, M.E., Crocetta, F., Golani, D., Rosso, A. et al., 2017. Uncertainties and validation of alien species catalogues: The Mediterranean as an example. Estuarine, Coastal and Shelf Science, 191, 171–187. https://doi.org/10.1016/j.ecss.2017.03.031

