## Supplementary material for "Species delimitation of eight ascidian (Tunicata) species from the North Eastern Mediterranean": Karahan_Supplementary data

Table S1. Sampling sites and date details

| Sites | Name of sites | Coordinates | *Botrylloides niger/ leachii | *Botrylloides israeliense | *Botrylloides sp. | *Botryllus schlosseri | *Botrylloides anceps | #Polyclinum constellatum | *Didemnum perlucidum | *Symplegma brakenhielmi |
|---|---|---|---|---|---|---|---|---|---|---|
| 1 | ♦Antalya-Kemer | 36.599871°N 30.575208°E | + | | Δ+ | | | | + | |
| 2 | ♦Antalya-Alanya | 36.559481°N 31.951682°E | + | | | | + | | +Δ | |
| 3 | ◊Mersin-Tisan | 36.157770°N 33.683355°E | + | +Δ | | | | | + | |
| 4 | ♦Mersin-Kızkalesi | 36.457684°N 34.144009°E | +Δ | +Δ | | | | +Δ | + | |
| 5 | ◊Mersin-Mezitli | 36.732885°N 34.521313°E | + | + | | | + | | + | +Δ |
| 6 | ◊Hatay-Konacık | 36.360380°N 35.820282°E | + | | | +Δ | +Δ | | + | |

* Sampling date-October 2018, # Sampling date-April 2014, ♦Stony bottom, ◊ Sandy bottom, Δ Bold uploaded specimen, + recorded in our censuses

Table S2. General information about specimen

| **Species name** | **N1** | **N2** | **BOLD ID and BIN ID** | **All specimen colors** | **Zooid distribution** |
|---|---|---|---|---|---|
| *Botrylloides niger/ aff. leachii* | 221 | 1 | IMS254-19, BOLD:ACI1328 | Diverse colors: from very light (creamy) to total black | 'leachii type' systems |
| *Botrylloides israeliense* | 36 | 2 | IMS260-20, BOLD:AEF0161<br><br>IMS261-20, BOLD:AEE6215 | Diverse colors: from light to dark black | 'leachii type' systems |
| *Botrylloides sp.* | 1 | 1 | IMS269-22, BOLD:AEC6032 | Brownish with the yellow strip | 'leachii type' systems in field, star shape in the aquaculture room |
| *Botryllus schlosseri* | 4 | 4 | IMS265-20, BOLD:AEE3528 | Brownish with a yellow strip, grey with white and yellow strip | Star shape |
| *Botrylloides anceps* | 27 | 1 | IMS259-20, BOLD:AEE3749 | Red, pink, creamy, brown, purple | 'leachii type' systems |
| *Symplegma brakenhielmi* | 1 | 1 | IMS264-20, BOLD:ADX5608 | Red | Zooids were randomly arranged |

| | | | | | |
|---|---|---|---|---|---|
| *Polyclinum constellatum* | 16 | 16 | IMS266-20, BOLD:ADC2025 | Light brown, reddish, and grey | Umbrella-like three-dimensional structure |
| *Didemnum perlucidum* | 2 | 2 | IMS267-20, BOLD:ACB6312 | White, orange, light and dark creamy, and brownish | Hard extracellular matrix, encrusted to the substratum |

N1: total sequenced sample number, N2: sample number used for species delimitation analysis

Table S3. Kimura-2 Parameter distance for bOTU-1 and -7.

| | Species name | OTU | 1 | 2 | 3 | 4 | 5 | 6 | 7 |
|---|---|---|---|---|---|---|---|---|---|
| 1 | *Botrylloides niger/aff.leachii* Kızkalesi (BN1) | 1 | | 0,00 | 0,00 | 0,00 | 0,00 | 0,00 | 0,03 |
| 2 | MT637961.1 *Botrylloides niger* | 1 | 0,00 | | 0,00 | 0,00 | 0,00 | 0,00 | 0,03 |
| 3 | MT637960.1 *Botrylloides niger* | 1 | 0,00 | 0,00 | | 0,00 | 0,00 | 0,00 | 0,03 |
| 4 | KU711783.1 *Botrylloides nigrum* | 1 | 0,01 | 0,01 | 0,01 | | 0,00 | 0,01 | 0,03 |
| 5 | LR828514.1 *Botrylloides niger* | 1 | 0,01 | 0,01 | 0,01 | 0,00 | | 0,01 | 0,03 |
| 6 | MG009579.1 *Botrylloides aff. leachii* | 1 | 0,00 | 0,00 | 0,00 | 0,01 | 0,01 | | 0,03 |
| 7 | *KF309549.1 Botrylloides leachii* | 6 | 0,19 | 0,19 | 0,19 | 0,19 | 0,19 | 0,19 | |

Above diagonal represents standard deviation. The red color refers to the present study sample and site, and the code in the brackets is the sample code.

Table S4. Kimura-2 Parameter distance for bOTU-13 and -12.

| | Species name | OTU | 1 | 2 | 3 | 4 | 5 | 6 |
|---|---|---|---|---|---|---|---|---|
| 1 | *Botrylloides israeliense* Tisan (G3) | 13 | | 0,01 | 0,01 | 0,01 | 0,01 | 0,03 |
| 2 | MG009580.1 *Botrylloides israeliense* | 13 | 0,03 | | 0,01 | 0,01 | 0,02 | 0,03 |
| 3 | MN076468.1 *Botrylloides sp.* | 13 | 0,03 | 0,02 | | 0,01 | 0,01 | 0,03 |
| 4 | *Botrylloides israeliense* Kızkalesi (Bts) | 13 | 0,05 | 0,05 | 0,04 | | 0,02 | 0,03 |
| 5 | MK978805.1 *Botrylloides sp.* | 13 | 0,07 | 0,08 | 0,07 | 0,08 | | 0,03 |
| 6 | MT873571.1 *Botrylloides sp.* | 12 | 0,18 | 0,19 | 0.18 | 0,18 | 0,16 | |

Above diagonal represents standard deviation. The red colors refer to the present study samples and sites, and the codes in the brackets is the sample codes.

Table S5. Kimura-2 Parameter distance for bOTU-16, -17, -18, -19 and -20.

| | Species name | OTU | 1 | 2 | 3 | 4 | 5 | 6 | 7 | 8 |
|---|---|---|---|---|---|---|---|---|---|---|
| 1 | MK575752. *Botryllus schlosseri* Clade A | 16 | | 0,010 | 0,010 | 0,012 | 0,019 | 0,020 | 0,025 | 0,028 |
| 2 | *Botryllus schlosseri* Konacık (L12) | 16 | 0,035 | | 0,005 | 0,008 | 0,018 | 0,019 | 0,023 | 0,025 |
| 3 | *Botryllus schlosseri* Konacık (L13) | 16 | 0,040 | 0,012 | | 0,007 | 0,018 | 0,019 | 0,024 | 0,027 |
| 4 | *Botryllus schlosseri* Konacık (L14) | 16 | 0,052 | 0,028 | 0,022 | | 0,020 | 0,022 | 0,026 | 0,029 |
| 5 | DQ340219. *Botryllus schlosseri* Clade B | 17 | 0,112 | 0,102 | 0,107 | 0,128 | | 0,017 | 0,023 | 0,022 |
| 6 | MK575788. *Botryllus schlosseri* Clade D | 18 | 0,126 | 0,110 | 0,115 | 0,136 | 0,095 | | 0,023 | 0,023 |
| 7 | DQ340214. *Botryllus schlosseri* Clade C | 19 | 0,163 | 0,145 | 0,151 | 0,173 | 0,139 | 0,141 | | 0,028 |
| 8 | MK575801. *Botryllus schlosseri* Clade E | 20 | 0,176 | 0,158 | 0,171 | 0,195 | 0,136 | 0,137 | 0,179 | |

Above diagonal represents standard deviation. The red colors refer to the present study samples and sites, and the codes in the brackets is the sample codes.

Table S6. Kimura-2 Parameter distance for bOTU-21 and -22

| | Species name | OTU | 1 | 2 | 3 | 4 |
|---|---|---|---|---|---|---|
| 1 | *Botrylloides anceps* Konacık (L8) | 21 | | 0,00 | 0,00 | 0,03 |
| 2 | MT637977. Botrylloides *sp.* | 21 | 0,00 | | 0,00 | 0,03 |
| 3 | MG009581. *Botrylloides anceps* | 21 | 0,01 | 0,01 | | 0,03 |
| 4 | MT873573. *Botrylloides cf. anceps* | 22 | 0,18 | 0,18 | 0,17 | |

Above diagonal represents standard deviation. The red color refers to the present study sample and site, and the code in the brackets is the sample code.

Table S7. Kimura-2 Parameter distance for bOTU-8 and -9.

| | Species name | OTU | 1 | 2 | 3 | 4 |
|---|---|---|---|---|---|---|
| 1 | *Botrylloides sp.* Kemer (C64) | 8 | | 0,003 | 0,003 | 0,02 |
| 2 | ON053356. *Botrylloides_sp.* | 8 | 0,00 | | 0,003 | 0,02 |
| 3 | ON053355. *Botrylloides_sp.* | 8 | 0,00 | 0,004 | | 0,02 |
| 4 | MT873575. *Botrylloides cf. pannosum* | 9 | 0,14 | 0,14 | 0,14 | |

Above diagonal represents standard deviation. The red colors refer to the present study samples and sites, and the codes in the brackets is the sample codes.

Table S8. Kimura-2 Parameter distance for bOTU-27 and -30.

| | Species name | OTU | 1 | 2 | 3 | 4 | 5 |
|---|---|---|---|---|---|---|---|
| 1 | *Symplegma brakenhielmi* Mezitli (Sp) | 30 | | 0,00 | 0,00 | 0,00 | 0,01 |
| 2 | MT232734.1 *Symplegma brakenhielmi* | 30 | 0,03 | | 0,00 | 0,00 | 0,01 |
| 3 | KX650765.1 *Botrylloides chevalense* | 30 | 0,04 | 0,01 | | 0,00 | 0,01 |
| 4 | KX650764.1 *Botrylloides chevalense* | 30 | 0,05 | 0,03 | 0,01 | | 0,01 |
| 5 | KU360802.1 *Symplegma brakenhielmi* | 27 | 0,29 | 0,29 | 0,29 | 0,27 | |

Above diagonal represents standard deviation. The red color refers to the present study sample and site, and the code in the brackets is the sample code.

Table S9. Kimura-2 Parameter distance for pOTU-1. Above diagonal represents standard deviation. The red colors refer to the present study samples and sites, and the codes in the brackets is the sample codes.

| | Species name | 1 | 2 | 3 | 4 | 5 | 6 | 7 | 8 | 9 | 10 | 11 | 12 | 13 | 14 | 15 | 16 | 17 | 18 | 19 | 20 | 21 | 22 | 23 | 24 |
|---|---|---|---|---|---|---|---|---|---|---|---|---|---|---|---|---|---|---|---|---|---|---|---|---|---|
| 1 | *Polyclinum constellatum* (1) Kızkalesi | | 0,00 | 0,00 | 0,00 | 0,00 | 0,00 | 0,00 | 0,00 | 0,00 | 0,00 | 0,00 | 0,00 | 0,00 | 0,00 | 0,00 | 0,00 | 0,00 | 0,00 | 0,00 | 0,00 | 0,01 | 0,01 | 0,01 | 0,01 |
| 2 | *Polyclinum constellatum* (2) Kızkalesi | 0,00 | | 0,00 | 0,00 | 0,00 | 0,00 | 0,00 | 0,00 | 0,00 | 0,00 | 0,00 | 0,00 | 0,00 | 0,00 | 0,00 | 0,00 | 0,00 | 0,00 | 0,00 | 0,00 | 0,01 | 0,01 | 0,01 | 0,01 |
| 3 | KX650781. *Polyclinum indicum* | 0,00 | 0,00 | | 0,00 | 0,00 | 0,00 | 0,00 | 0,00 | 0,00 | 0,00 | 0,00 | 0,00 | 0,00 | 0,00 | 0,00 | 0,00 | 0,00 | 0,00 | 0,00 | 0,00 | 0,01 | 0,01 | 0,01 | 0,01 |
| 4 | MT873559. *Polyclinum constellatum* | 0,00 | 0,00 | 0,00 | | 0,00 | 0,00 | 0,00 | 0,00 | 0,00 | 0,00 | 0,00 | 0,00 | 0,00 | 0,00 | 0,00 | 0,00 | 0,00 | 0,00 | 0,00 | 0,00 | 0,01 | 0,01 | 0,01 | 0,01 |
| 5 | *Polyclinum constellatum* (3) Kızkalesi | 0,00 | 0,00 | 0,00 | 0,00 | | 0,00 | 0,00 | 0,00 | 0,00 | 0,00 | 0,00 | 0,00 | 0,00 | 0,00 | 0,00 | 0,00 | 0,00 | 0,00 | 0,00 | 0,00 | 0,01 | 0,01 | 0,01 | 0,01 |
| 6 | *Polyclinum constellatum* (4) Kızkalesi | 0,00 | 0,00 | 0,00 | 0,00 | 0,00 | | 0,00 | 0,00 | 0,00 | 0,00 | 0,00 | 0,00 | 0,01 | 0,00 | 0,00 | 0,00 | 0,00 | 0,00 | 0,00 | 0,00 | 0,01 | 0,01 | 0,01 | 0,01 |
| 7 | OL597608. *Polyclinum constellatum* | 0,01 | 0,01 | 0,01 | 0,01 | 0,01 | 0,00 | | 0,00 | 0,00 | 0,00 | 0,01 | 0,01 | 0,01 | 0,00 | 0,00 | 0,00 | 0,00 | 0,00 | 0,00 | 0,01 | 0,01 | 0,01 | 0,01 | 0,01 |
| 8 | MW285146.1 *Polyclinum constellatum* | 0,01 | 0,01 | 0,01 | 0,01 | 0,01 | 0,00 | 0,00 | | 0,00 | 0,00 | 0,01 | 0,01 | 0,01 | 0,00 | 0,00 | 0,00 | 0,00 | 0,00 | 0,00 | 0,01 | 0,01 | 0,01 | 0,01 | 0,01 |
| 9 | KX650778. *Polyclinum indicum* | 0,00 | 0,00 | 0,00 | 0,00 | 0,00 | 0,00 | 0,01 | 0,01 | | 0,00 | 0,00 | 0,00 | 0,01 | 0,00 | 0,00 | 0,00 | 0,00 | 0,00 | 0,00 | 0,00 | 0,01 | 0,01 | 0,01 | 0,01 |
| 10 | KX650773. *Polyclinum indicum* | 0,01 | 0,01 | 0,01 | 0,01 | 0,01 | 0,00 | 0,01 | 0,01 | 0,01 | | 0,01 | 0,01 | 0,01 | 0,00 | 0,00 | 0,00 | 0,00 | 0,00 | 0,01 | 0,01 | 0,01 | 0,01 | 0,01 | 0,01 |
| 11 | KX650779. *Polyclinum indicum* | 0,01 | 0,01 | 0,01 | 0,01 | 0,01 | 0,01 | 0,02 | 0,02 | 0,01 | 0,01 | | 0,01 | 0,01 | 0,00 | 0,00 | 0,00 | 0,00 | 0,01 | 0,01 | 0,01 | 0,01 | 0,01 | 0,01 | 0,01 |
| 12 | *Polyclinum constellatum* (5) Kızkalesi | 0,01 | 0,01 | 0,01 | 0,01 | 0,01 | 0,01 | 0,02 | 0,02 | 0,01 | 0,02 | 0,02 | | 0,00 | 0,00 | 0,00 | 0,00 | 0,00 | 0,00 | 0,00 | 0,00 | 0,00 | 0,01 | 0,01 | 0,01 |
| 13 | *Polyclinum constellatum* (6) Kızkalesi | 0,01 | 0,01 | 0,01 | 0,01 | 0,01 | 0,01 | 0,02 | 0,02 | 0,01 | 0,02 | 0,02 | 0,01 | | 0,00 | 0,00 | 0,00 | 0,00 | 0,00 | 0,01 | 0,01 | 0,01 | 0,01 | 0,01 | 0,01 |
| 14 | *Polyclinum constellatum* (7) Kızkalesi | 0,00 | 0,00 | 0,00 | 0,00 | 0,00 | 0,00 | 0,01 | 0,01 | 0,00 | 0,01 | 0,01 | 0,01 | 0,01 | | 0,00 | 0,00 | 0,00 | 0,00 | 0,00 | 0,00 | 0,01 | 0,01 | 0,01 | 0,01 |
| 15 | *Polyclinum constellatum* (8) Kızkalesi | 0,00 | 0,00 | 0,00 | 0,00 | 0,00 | 0,00 | 0,01 | 0,01 | 0,00 | 0,01 | 0,01 | 0,01 | 0,01 | 0,00 | | 0,00 | 0,00 | 0,00 | 0,00 | 0,00 | 0,01 | 0,01 | 0,01 | 0,01 |
| 16 | *Polyclinum constellatum* (9) Kızkalesi | 0,00 | 0,00 | 0,00 | 0,00 | 0,00 | 0,00 | 0,01 | 0,01 | 0,00 | 0,01 | 0,01 | 0,01 | 0,01 | 0,00 | 0,00 | | 0,00 | 0,00 | 0,00 | 0,00 | 0,01 | 0,01 | 0,01 | 0,01 |
| 17 | *Polyclinum constellatum* (10) Kızkalesi | 0,00 | 0,00 | 0,00 | 0,00 | 0,00 | 0,00 | 0,01 | 0,01 | 0,00 | 0,01 | 0,01 | 0,01 | 0,01 | 0,00 | 0,00 | 0,00 | | 0,00 | 0,00 | 0,00 | 0,01 | 0,01 | 0,01 | 0,01 |
| 18 | *Polyclinum constellatum* (11) Kızkalesi | 0,00 | 0,00 | 0,00 | 0,00 | 0,00 | 0,00 | 0,01 | 0,01 | 0,01 | 0,01 | 0,01 | 0,01 | 0,01 | 0,00 | 0,00 | 0,00 | 0,00 | | 0,00 | 0,00 | 0,01 | 0,01 | 0,01 | 0,01 |
| 19 | *Polyclinum constellatum* (12) Kızkalesi | 0,01 | 0,01 | 0,01 | 0,01 | 0,01 | 0,01 | 0,01 | 0,01 | 0,01 | 0,01 | 0,02 | 0,01 | 0,02 | 0,01 | 0,01 | 0,01 | 0,01 | 0,01 | | 0,00 | 0,00 | 0,01 | 0,01 | 0,01 |
| 20 | *Polyclinum constellatum* (13) Kızkalesi | 0,01 | 0,01 | 0,01 | 0,01 | 0,01 | 0,01 | 0,01 | 0,01 | 0,01 | 0,02 | 0,02 | 0,00 | 0,01 | 0,01 | 0,01 | 0,01 | 0,01 | 0,01 | 0,00 | | 0,00 | 0,01 | 0,01 | 0,01 |
| 21 | *Polyclinum* constellatum (14) Kızkalesi | 0,02 | 0,02 | 0,02 | 0,02 | 0,02 | 0,02 | 0,02 | 0,02 | 0,02 | 0,03 | 0,03 | 0,01 | 0,02 | 0,02 | 0,02 | 0,02 | 0,02 | 0,02 | 0,01 | 0,01 | | 0,01 | 0,01 | 0,01 |
| 22 | *Polyclinum constellatum* (15) Kızkalesi | 0,02 | 0,02 | 0,02 | 0,02 | 0,02 | 0,02 | 0,03 | 0,03 | 0,03 | 0,03 | 0,04 | 0,02 | 0,03 | 0,02 | 0,02 | 0,02 | 0,02 | 0,02 | 0,02 | 0,03 | 0,03 | | 0,01 | 0,01 |
| 23 | *Polyclinum constellatum* (16) Kızkalesi | 0,03 | 0,03 | 0,03 | 0,03 | 0,03 | 0,03 | 0,04 | 0,04 | 0,03 | 0,04 | 0,03 | 0,04 | 0,04 | 0,03 | 0,03 | 0,03 | 0,03 | 0,03 | 0,04 | 0,04 | 0,05 | 0,05 | | 0,01 |
| 24 | KX138512. *Polyclinum sp.* | 0,05 | 0,05 | 0,05 | 0,05 | 0,05 | 0,05 | 0,06 | 0,06 | 0,05 | 0,07 | 0,07 | 0,06 | 0,06 | 0,05 | 0,05 | 0,05 | 0,05 | 0,05 | 0,06 | 0,05 | 0,06 | 0,07 | 0,08 | |

Table S10. Kimura-2 Parameter distance for dOTU-1

| | Species name | 1 | 2 | 3 | 4 | 5 | 6 | 7 | 8 | 9 | 10 | 11 | 12 |
|---|---|---|---|---|---|---|---|---|---|---|---|---|---|
| 1 | MH824680 *Didemnum perlucidum* | | 0,003 | 0,004 | 0,006 | 0,000 | 0,000 | 0,003 | 0,004 | 0,003 | 0,003 | 0,003 | 0,012 |
| 2 | KX650791 *Didemnum perlucidum* | 0,006 | | 0,002 | 0,006 | 0,003 | 0,003 | 0,000 | 0,002 | 0,000 | 0,000 | 0,000 | 0,011 |
| 3 | KU883151 *Didemnum perlucidum* | 0,008 | 0,002 | | 0,006 | 0,004 | 0,004 | 0,002 | 0,002 | 0,002 | 0,002 | 0,002 | 0,012 |
| 4 | KX138490 *Didemnum perlucidum* | 0,022 | 0,016 | 0,016 | | 0,006 | 0,006 | 0,006 | 0,006 | 0,006 | 0,006 | 0,006 | 0,014 |
| 5 | KU667270 *Didemnum perlucidum* | 0,000 | 0,006 | 0,008 | 0,022 | | 0,000 | 0,003 | 0,004 | 0,003 | 0,003 | 0,003 | 0,012 |
| 6 | KR537439 *Didemnum perlucidum* | 0,000 | 0,006 | 0,008 | 0,022 | 0,000 | | 0,003 | 0,004 | 0,003 | 0,003 | 0,003 | 0,012 |
| 7 | JQ731735. *Didemnum perlucidum* | 0,006 | 0,000 | 0,002 | 0,016 | 0,006 | 0,006 | | 0,002 | 0,000 | 0,000 | 0,000 | 0,011 |
| 8 | MT637962 *Didemnum perlucidum* | 0,008 | 0,002 | 0,002 | 0,016 | 0,008 | 0,008 | 0,002 | | 0,002 | 0,002 | 0,002 | 0,012 |
| 9 | MN184710 *Didemnum perllucidum*-Australia | 0,006 | 0,000 | 0,002 | 0,016 | 0,006 | 0,006 | 0,000 | 0,002 | | 0,000 | 0,000 | 0,011 |
| 10 | *Didemnum perlucidum* (C2 ) | 0,006 | 0,000 | 0,002 | 0,016 | 0,006 | 0,006 | 0,000 | 0,002 | 0,000 | | 0,000 | 0,011 |
| 11 | *Didemnum perlucidum* (C3) | 0,006 | 0,000 | 0,002 | 0,016 | 0,006 | 0,006 | 0,000 | 0,002 | 0,000 | 0,000 | | 0,011 |
| 12 | KY741541 *Didemnum etiolum* | 0,064 | 0,057 | 0,059 | 0,076 | 0,064 | 0,064 | 0,057 | 0,059 | 0,057 | 0,057 | 0,057 | |

Above diagonal represents standard deviation. The red colors refer to the present study samples and sites, and the codes in the brackets is the sample codes.

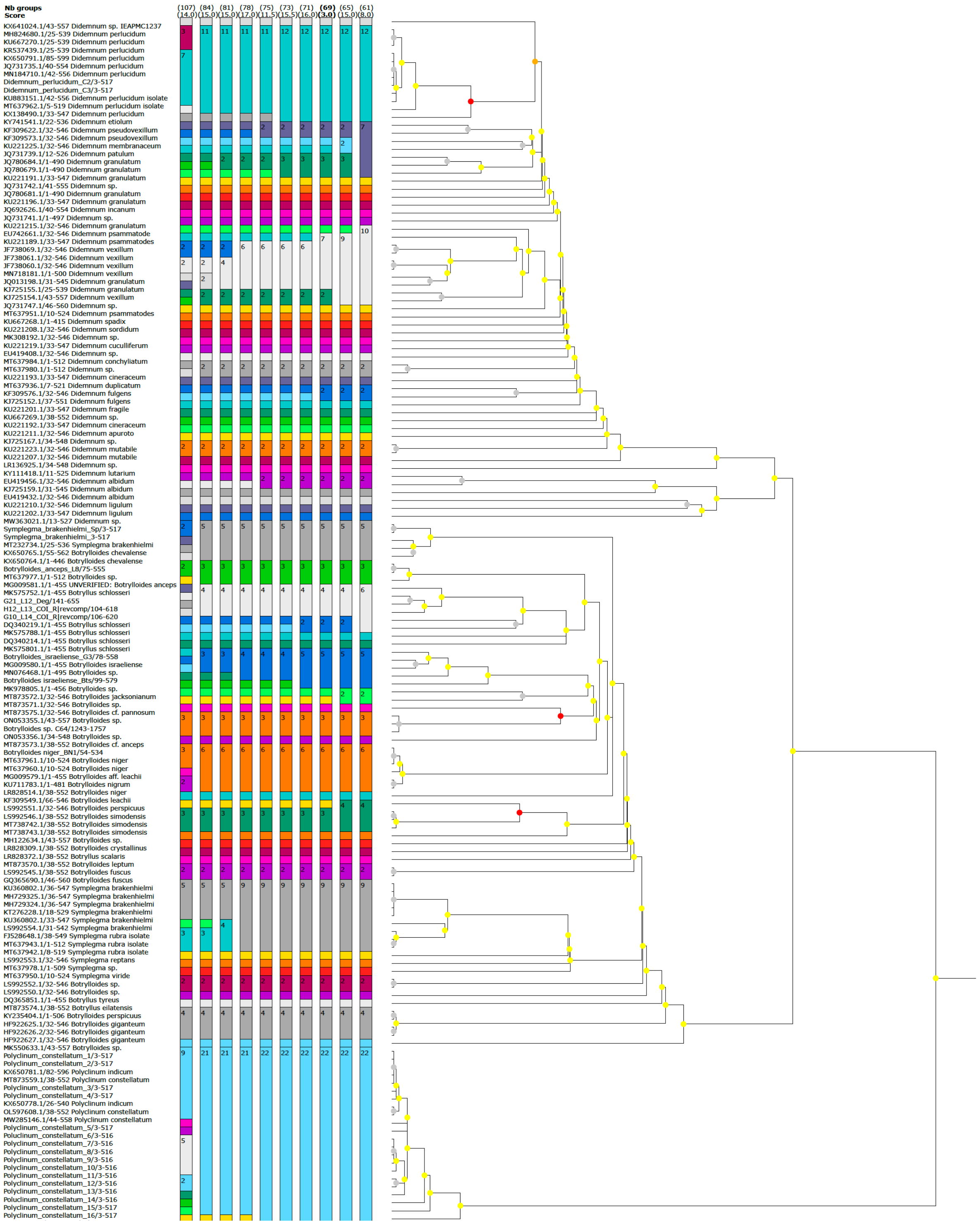


Fig. S1. ASAP score; Colors represent different OTUs. Number in line of the OTUs present the total samples assigned to the same OTU. The first number line above the OTUs' columns presents the total OTU numbers, values at the second line indicate ASAP-scores (the lowest the score the better is the partition, Puillandre et al. 2021).

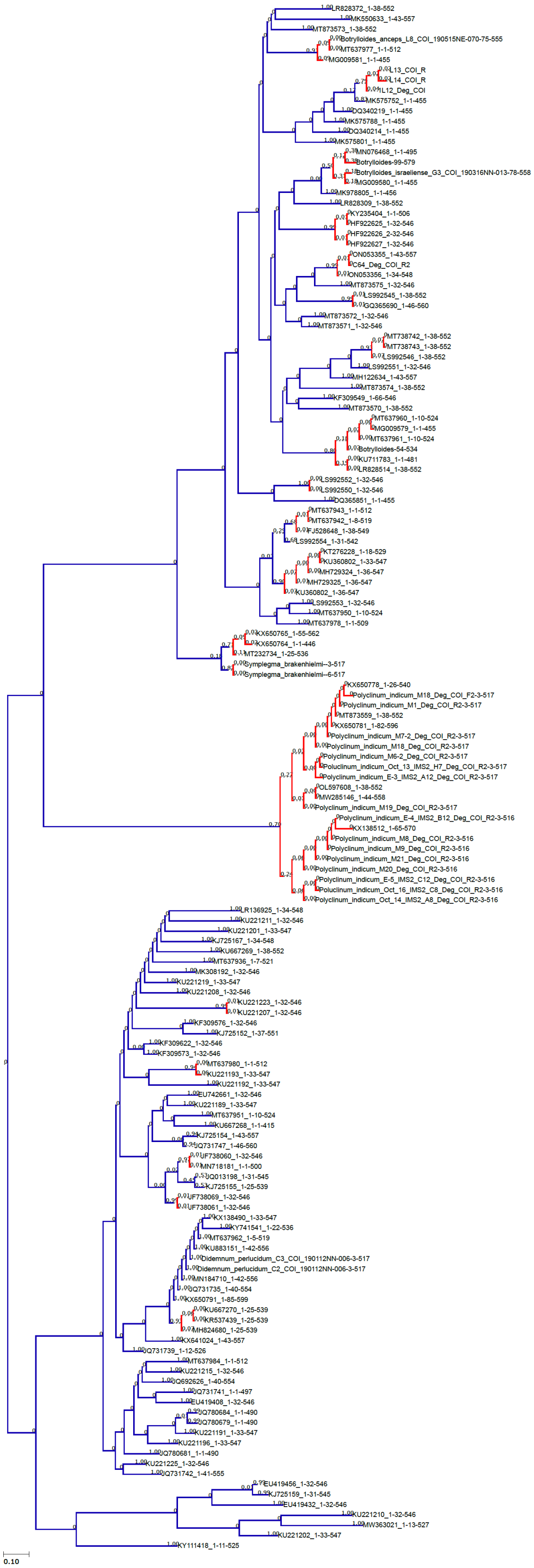


Fig. S2. PTP score; Blue lines present different OTUs, red lines present same OTUs

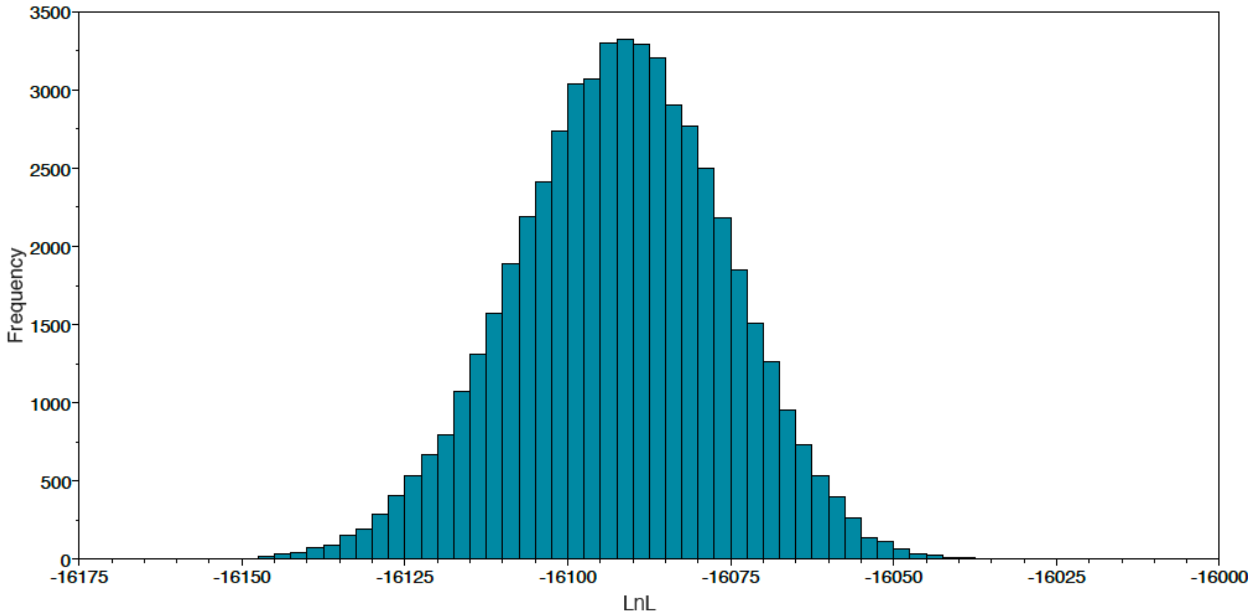


Fig. S3. LnL value of MrBayes analysis.

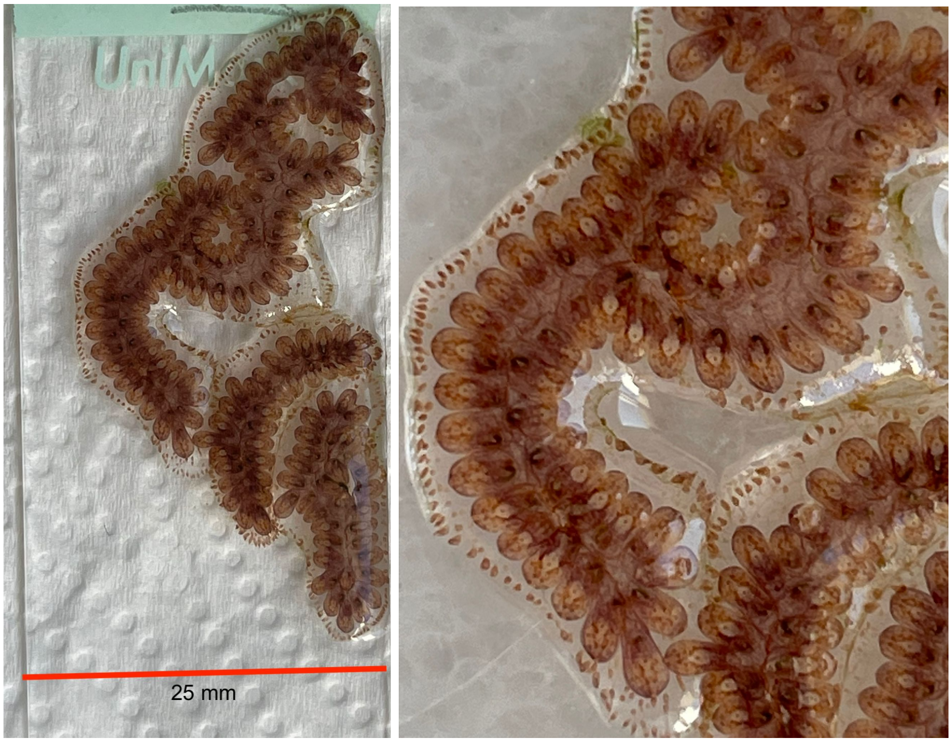


Fig. S4. When the zooid size of the brown morphotypes of *B. anceps* reached up to 2 mm (the photograps were captured via a cell phone).

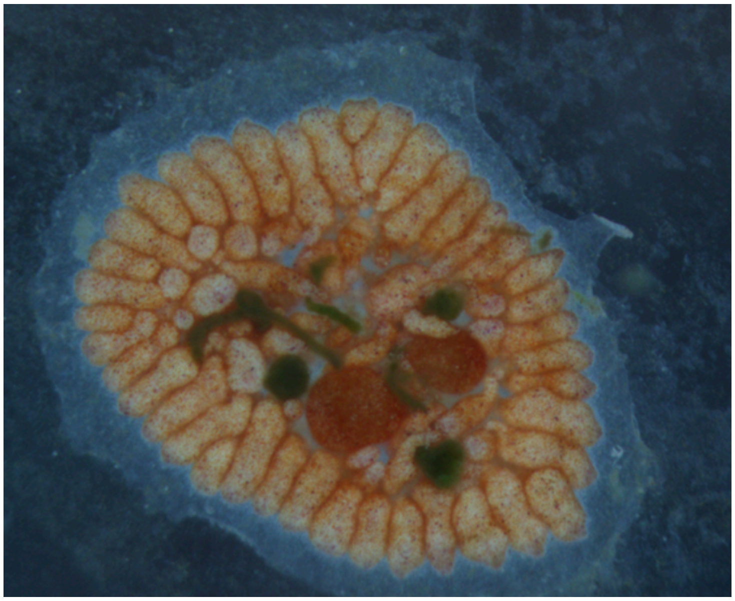

Fig. S5. Large, retracted and condensed ampullas of *Botrylloides anceps*

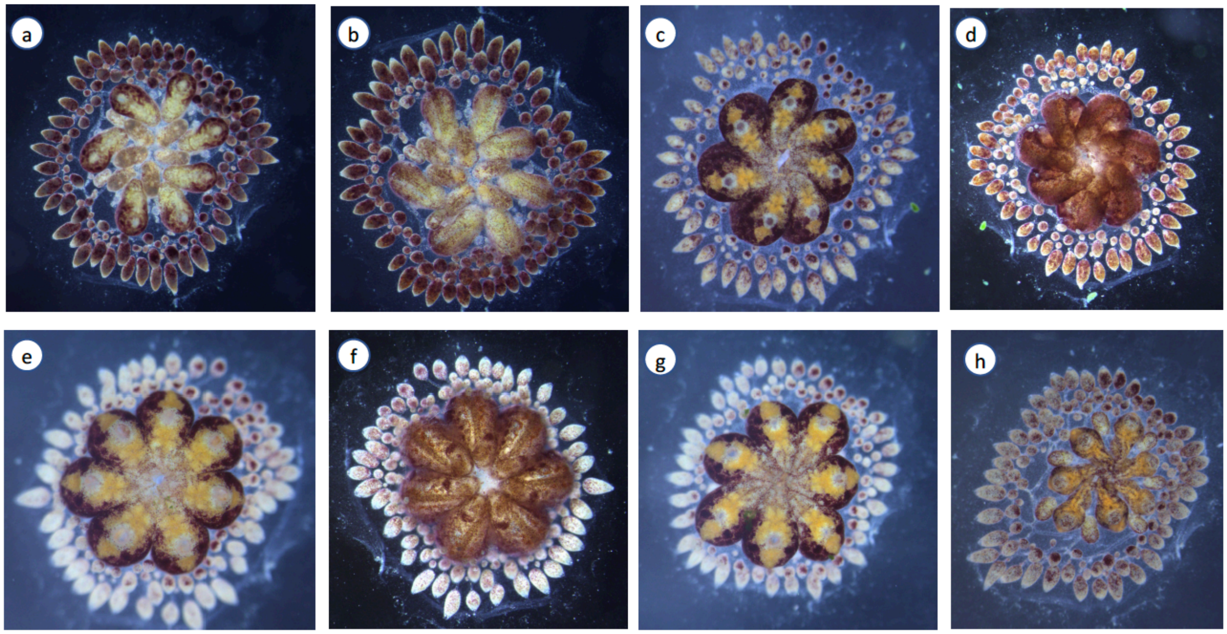


Fig. S6. Blastogenic cycle of *Botrylloides sp.* Dorsal and ventral view of the Blastogenic stages; a-b) stage-D (Jun 07, 2018), c-d) stage-A (Jun 08, 2018), e-f) stage-B (Jun 09, 2018), g) Dorsal-stage C (Jun 10, 2018), h) Dorsal – stage D (Jun 11, 2018). Abbreviations: end; endostyle, st; stomach, cco; common cloacal opening, sg: stigmata, a: ampullas.
